# Old World alphaviruses use distinct mechanisms to infect brain microvascular endothelial cells for neuroinvasion

**DOI:** 10.1101/2025.01.22.634395

**Authors:** Pablo A. Alvarez, Ashley Tang, Declan M. Winters, Prashant Kaushal, Angelica Medina, Karolina E. Kaczor-Urbanowicz, Bryan Ramirez Reyes, Robyn M. Kaake, Oliver I. Fregoso, April D. Pyle, Mehdi Bouhaddou, Hengli Tang, Melody M.H. Li

## Abstract

Several alphaviruses bypass the blood-brain barrier (BBB), causing debilitating or fatal encephalitis. Sindbis virus (SINV) has been extensively studied *in vivo* to understand alphavirus neuropathogenesis; yet the molecular details of neuroinvasion at the BBB remain poorly understood. We investigated alphavirus-BBB interactions by pairing a physiologically relevant, human pluripotent stem cell derived model of brain microvascular endothelial cells (BMECs) with SINV strains of opposite neuroinvasiveness. Our system demonstrates that SINV neuroinvasion correlates with robust infection of the BBB. Specifically, SINV genetic determinants of neuroinvasion enhance viral entry into BMECs. We also identify solute carrier family 2 member 3 (SLC2A3, also named GLUT3) as a potential BMEC-specific entry factor exploited for neuroinvasion. Strikingly, efficient BBB infection is a conserved phenotype that correlates with the neuroinvasive capacity of several Old World alphaviruses, including chikungunya virus. Here, we reveal BBB infection as a shared pathway for alphavirus neuroinvasion that can be targeted for preventing alphavirus-induced encephalitis.

## INTRODUCTION

Alphavirus-induced neuropathies pose a significant global threat to pediatric, geriatric, and immunocompromised populations. Among the alphaviruses, Eastern equine encephalitis virus (EEEV) and chikungunya virus (CHIKV) stand out as the deadliest and most widespread, respectively, both capable of causing severe neurological disorders in humans. While EEEV cases are rare, up to 50% of infections in the Northeastern United States result in fatal encephalitis^1^. The more widespread CHIKV is primarily studied for its arthritogenic effects; however, the largest recorded outbreak on La Reunion island resulted in 60-80% of patients experiencing neurological complications, including seizures and encephalitis^2,3^. Notably, recent findings identified CHIKV RNA in the cerebrospinal fluid of all fatal cases within a human cohort, underscoring the critical role of neuroinvasion in disease outcomes^4^. To establish infection in the central nervous system (CNS), alphaviruses rely on hematogenous dissemination and must bypass the blood-brain barrier (BBB)^5^. Consequently, elucidating the molecular alphavirus-BBB interactions driving neuroinvasion is essential for developing countermeasures against alphavirus-induced neuropathies.

The BBB is the primary obstacle preventing alphaviruses from accessing the CNS^6–8^. It is composed of brain microvascular endothelial cells (BMECs) structurally supported by pericytes and astrocytes^9^. Crossing the BBB can occur through three primary mechanisms: 1) transcellular transport through BMECs, 2) paracellular migration between BMECs, or 3) transport by immune cells such as macrophages^10,11^. However, mounting evidence suggests that alphaviruses do not rely on barrier disruption or immune cell-mediated carriage to cross the BBB^12–14^. Studies on Venezuelan equine encephalitis virus (VEEV) suggests that it exploits vesicle trafficking machinery in BMECs to achieve neuroinvasion^12^. Additionally, VEEV and the related Western equine encephalitis virus (WEEV) were shown to infect BMECs, pericytes, and astrocytes; however, whether these infections contribute directly to neuroinvasion remains unclear.

Recent studies show that human pluripotent stem cell (hPSC) derived brain microvascular endothelial-like cells (hPSC-BMELCs) accurately recapitulate *in vivo* functionality of the BBB^15^. Additionally, hPSC-BMELCs share the distinct gene expression profiles of primary human BMECs^16^ and can be further driven towards endothelial maturation with the introduction of specific transcription factors^17,18^, making the former a physiologically relevant model for investigating molecular interactions at the BBB. This model has been employed to investigate host BBB-pathogen interactions for several microbes including *Plasmodium falciparum*, *Streptococcus agalactiae*, and SARS-CoV-2, that cause malaria, group B strep infections, and COVID-19, respectively^19–21^. For arboviruses, previous work demonstrated that hPSC-BMELCs recapitulate the neuroinvasive phenotypes of alphaviruses and flaviviruses observed *in vivo.* Moreover, this model was used to identify the interferon-stimulated gene IFITM1 that protects against flavivirus neuroinvasion^22^. Specifically, this model demonstrated that non-neuroinvasive flaviviruses are inhibited by IFITM1, while neuroinvasive flaviviruses have evolved mechanisms to bypass this restriction. Thus, this is a great model for identifying host factors that differentially promote or inhibit different viral strains, especially in the context of neuroinvasion.

Historically, research into alphavirus neuroinvasion has concentrated on New World alphaviruses. However, Old World alphaviruses such as Sindbis virus (SINV), Semliki Forest virus (SFV), and CHIKV also exhibit neurotropism in humans^2,23,24^. CHIKV remains particularly understudied in this context due to the lack of *in vivo* models that faithfully replicate its neurological complications, with macaques being a notable exception^25^. In contrast, SINV is widely used as a model for alphavirus neuroinvasion and pathogenesis because of the availability of robust mouse models and the ability to study it outside of biosafety level 3 containment. Over time, this accessibility has led to the development of valuable tools and strains for investigating SINV neuroinvasion. For instance, serial passage of a SINV isolate in the brains of young mice resulted in a neurovirulent strain that was not neuroinvasive, named SVN. SVN was further passaged to produce SVNI, which was both neurovirulent and neuroinvasive^6^. Subsequent work identified the genetic determinants of fatal SINV neuroinvasion in mice, culminating in the creation of the recombinant strain, R47^7^. R47 has three key differences from SVN: a mutation in the 5’ noncoding region that is homologous to the nucleotide changes in VEEV that confer resistance to the interferon-stimulated gene IFIT1^26^, and two amino acid changes in the E2 glycoprotein. While these mutations are critical for neuroinvasion, their mechanisms of action and functional consequences are not yet understood.

The alphavirus E2 glycoprotein is central to virus entry and assembly^5,27,28^. Critical residues in E2, such as residues 190 and 260 in SINV and residue 82 in CHIKV, have been identified as key determinants of neuroinvasion and pathogenicity *in vivo*^7,29^. During early stages of infection, E2 facilitates host cell attachment and receptor engagement, processes essential for viral entry^5^. Since alphaviruses infect various cell types in both vertebrate and invertebrate hosts, the E2 protein must interact with several different host attachment factors and receptors, collectively called entry factors^5,30^. These include glycosaminoglycans, such as heparan sulfate and chondroitin sulfate, which are commonly utilized for initial attachment^30^. For internalization and fusion, alphaviruses use various cell surface protein receptors in a lineage-specific manner: most Old World alphaviruses engage matrix remodeling-associated protein 8 (MXRA8) in a species specific manner, while New World alphaviruses rely on several low-density lipoprotein receptors, including very low-density lipoprotein receptor (VLDLR), low-density lipoprotein receptor (LDLR), low-density lipoprotein receptor class A domain containing 3 (LDLRAD3), and LDLR related protein 8 (LRP8, also named ApoER2)^30–35^. Other receptors, including protocadherin 10 (PCDH10) and solute carrier family 11 member 2 (SLC11A2, also named natural resistance-associated macrophage protein 2, or NRAMP2), have also been identified^36,37^. Despite these advances in receptor identification, the physiological relevance of cell type-specific expression of these entry factors and their contribution to alphavirus pathogenesis remains largely unexplored.

Here, we show that neuroinvasive ability of alphaviruses *in vivo* correlates with the ability to infect all cells of the BBB. Specifically, we demonstrate that neuroinvasive-specific residues in SINV enhance viral attachment, internalization, and fusion to BMECs, driving BBB infection and subsequent neuroinvasion. Our differential analysis of transcriptomic data across endothelial cells revealed that only a handful of known receptors are BMEC-specific; however, functional experiments suggest a novel entry factor is responsible for the differential infection phenotype in BMECs. Hence, we performed proteomics to identify highly expressed proteins on the cell surface that are unique to BMECs, and uncovered solute carrier family 2 member 3 (SLC2A3, also named glucose transporter 3 (GLUT3)) as a novel candidate receptor for R47 specific infection of BMECs. Notably, we also demonstrate that other Old World alphaviruses, including the clinically relevant CHIKV, infect BBB cells in a manner consistent with their neuroinvasive capacity *in vivo*. These findings provide crucial insights into the molecular interactions between alphaviruses and the host BBB and offer a foundation for the development of innovative countermeasures to prevent neuroinvasion and mitigate alphavirus-induced neuropathies.

## RESULTS

### Neuroinvasive SINV efficiently infects cells of the blood-brain barrier

The BBB is primarily composed of BMECs that are structurally supported by pericytes and astrocyte end feet^9^. Infection of these cell types is well documented for some alphaviruses and flaviviruses^12,38,39^, yet how infection of these cells contributes to neuroinvasion is less understood. Previous work demonstrated that the ability to infect hPSC-BMELCs is dependent on the neuroinvasive capacity of SINV^22^. To confirm that this is specific to endothelial cells in the brain, we investigated this differential infection phenotype across various endothelial cell models. We first generated hPSC-BMELCs, which express the proper endothelial cell markers such as platelet endothelial cell adhesion molecule 1 (PECAM-1, also named CD31), active transporters such as glucose transporter 1 (GLUT1), and tight junction proteins such as claudin 5 (CLDN-5)^16,40–42^ (Figure S1A-C). To ensure that the differential infection phenotype between SVN and R47 is not an artifact of differing virus input, we infected hPSC-BMELCs with various MOIs of SVN and R47. Significant differential infection was observed for all MOIs tested; however, the difference is lower at the highest MOI of 10, likely due to saturation (Figure S2A). We next tested multiple MOIs in human umbilical vein endothelial cells (HUVECs), primary human BMECs (HBMECs), and immortalized human BMECs (HCMEC/D3). As expected, HUVECs were efficiently infected by both strains at every MOI, suggesting that the ability to infect peripheral endothelial cells is conserved for both neuroinvasive and non-neuroinvasive SINV strains (Figure S2B). HBMECs were differentially infected at all MOIs tested but only statistically significant at an MOI of 10. Interestingly, we also observed a lower maximum infection rate in HBMECs —25% for R47 at an MOI of 10, compared to almost 90% in hPSC-BMELCs (Figure S2C). Surprisingly, HCMEC/D3 cells were not infected regardless of MOI (Figure S2D). This is likely due to key differences between hPSC-BMELCs and HCMEC/D3 cells, such as the low barrier integrity of HCMEC/D3 cells compared to that of HBMECs and hPSC-BMELCs as shown in our previous study^22^. Due to the conserved differential infection phenotype between hPSC-BMELCs and HBMECs, we proceeded with hPSC-BMELCs to thoroughly investigate the mechanism behind efficient infection of the BBB with R47.

To better understand BBB infection as a possible route of neuroinvasion for SINV, we compared the ability of these strains to infect all cells of the neurovascular unit. An infection time course in hPSC-BMELCs reveals that R47 has a 2-fold advantage over SVN as early as 12 hours post infection (Figure 1A). Strikingly, pericytes are also differentially infected by SVN and R47, albeit the rate of infection is slower, where R47 has an advantage over SVN beginning at 18 hours post infection (Figure 1B). Astrocyte infection revealed that both SVN and R47 replicate to high levels already by 18 hours post infection, even though SVN has a slight advantage over R47, which diminishes over time and is completely gone by 24 hours post infection (Figure 1C). We next decided to investigate virion production in each of these cell types. For both hPSC-BMELCs and pericytes, there is greater R47 virion production as early as 12 hours post infection, however the major difference between SVN and R47 does not occur until 24 hours post infection (Figure 1D and 1E). Additionally, the differential infection phenotype is amplified in hPSC-BMELCs, where previously the difference in percent infection is 2-3 fold (Figure 1A), now the difference in virion production is almost 1000-fold (Figure 1D). Strikingly, we also observed a 100-fold increase in R47 virion production compared to SVN in astrocytes, despite no advantage in viral replication indicated by GFP expression (Figure 1F). Together, these data demonstrate that neuroinvasive SINV has gained the ability to more efficiently replicate in BMECs and pericytes, and produce virus particles more efficiently in all cell types of the BBB.

**Figure 1.**
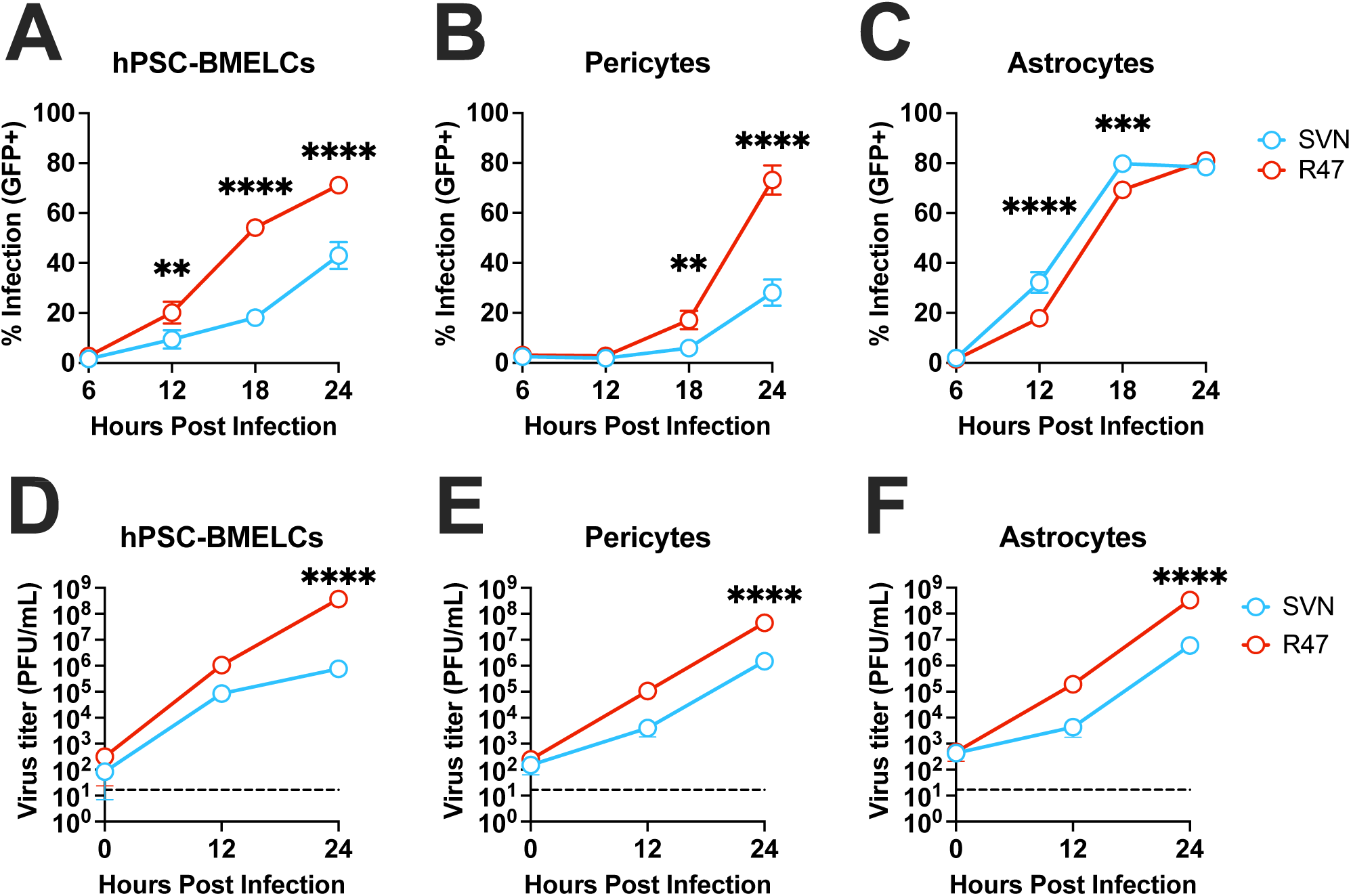
Neuroinvasive SINV efficiently infects all cells of the blood-brain barrier. (A) hPSC-BMELCs, (B) Pericytes, or (C) Astrocytes were infected with either GFP-expressing SVN or R47 at an MOI of 0.1 and harvested for flow cytometry at 6, 12, 18, or 24 hours post infection. (D) hPSC-BMELCs, (E) Pericytes, or (F) Astrocytes were infected with parental SVN or R47 viruses at an MOI of 0.1. Media was harvested at 0, 12, or 24 hours post infection for plaque forming assays using BHK-21 cells. Asterisks indicate statistically significant differences between SVN and R47 at the indicated time point (two-way ANOVA and Šidák’s multiple comparisons): **, *p* < 0.01; ***, *p* < 0.001; ****, *p* < 0.0001. Non-significant differences are not shown. Data are representative of 2 independent experiments. Data is shown as mean ±SD.

### BMEC infection and virion production precedes barrier breakdown

Work *in vivo* suggests that BBB disruption is a consequence of alphavirus neuroinvasion^13^. We chose to follow up on this work *in vitro* using the versatility of the hPSC-BMELC model. hPSC-BMELCs exhibit physiologically relevant barrier properties, and thus provide an excellent model for observing changes in barrier function *in vitro*^16,22^. To mimic the BBB *in vitro* we cultured hPSC-BMELCs in a transwell system, where hPSC-BMELCs are seeded in a transwell insert that separates a larger well into two distinct compartments. The inner compartment, referred to as the apical chamber, represents the inside of the blood vessels, made up of BMECs. The outer compartment, referred to as the basolateral chamber, represents the brain parenchyma outside of the blood vessels. Therefore, to investigate whether SINV breaks down the BBB and whether this happens prior to BMEC infection, we infected hPSC-BMELCs in the transwell system and measured barrier integrity via transendothelial electrical resistance (TEER) over time (Figure 2A). At the same time, we measured virus crossing by determining the amount of infectious virus production in the basolateral chamber at each time point via plaque assay. TEER values for hPSC-BMELCs began at around 1500Ωcm^2^, which is close to the expected value of human BMECs *in vivo*^15^. Barrier integrity then decreases by half for both SVN and R47 infected hPSC-BMELCs at 18 hours post infection; however, the TEER for SVN infected hPSC-BMELCs plateaus at about 600Ωcm^2^ while the TEER for R47 infected hPSC-BMELCs reaches almost 0Ωcm^2^ by 24 hours post infection (Figure 2B). Importantly, 100-fold more R47 is detected in the basolateral chamber compared to SVN as early as 12 hours post infection, and is consistent at 18 and 24 hours post infection (Figure 2C). Moreover, decreases in TEER remain the same for both SVN and R47 until 24 hours post infection. Therefore, decreases in TEER do not explain the differential detection of SVN and R47 virion particles early on in the basolateral chamber. Together, these data demonstrate barrier leakage to be a consequence of BMEC infection and support BMEC infection as a major route of SINV neuroinvasion.

**Figure 2.**
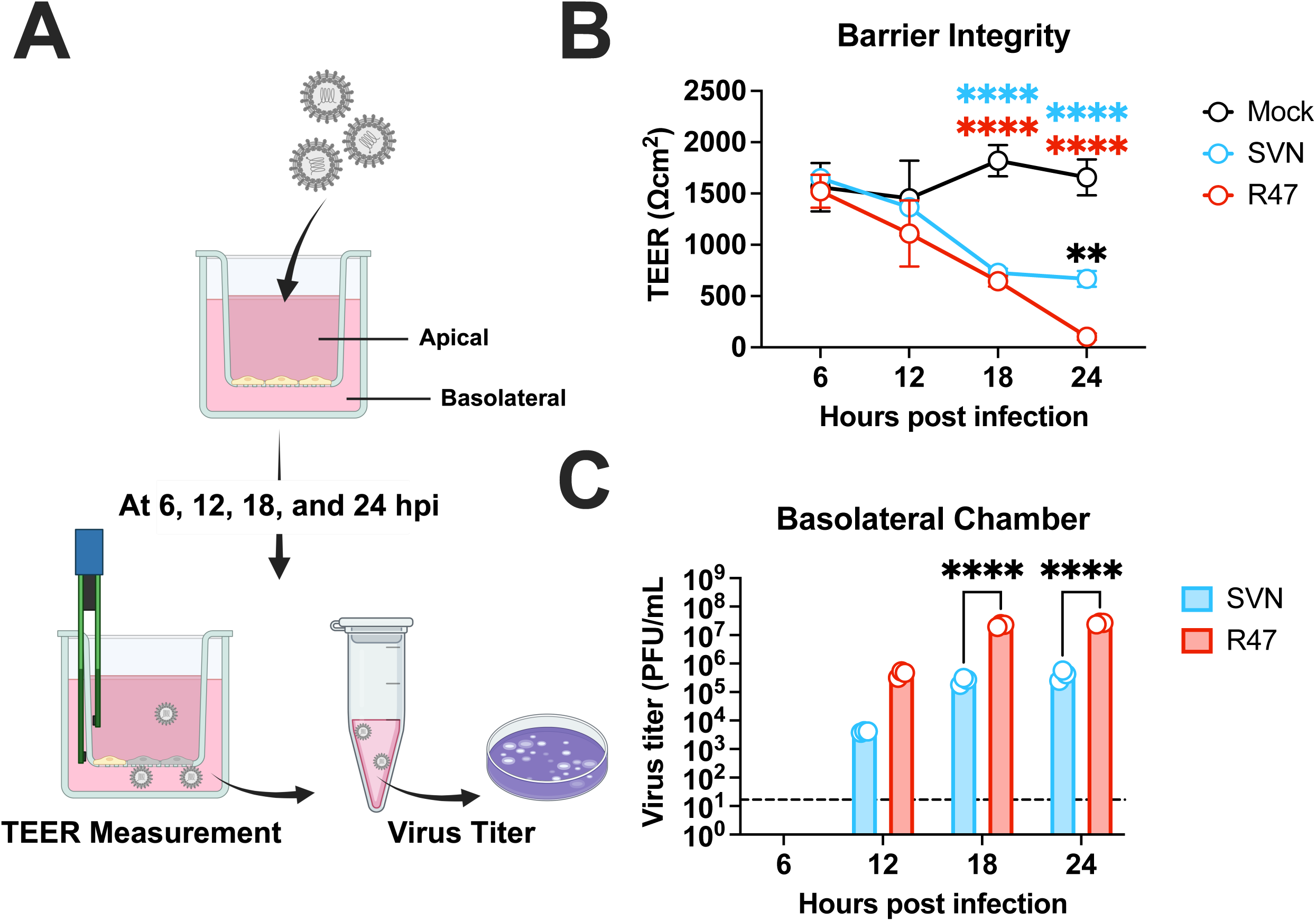
BMEC infection and virion production precedes barrier breakdown. (A) Experimental schematic. Briefly, hPSC-BMELCs were seeded in the apical side of a transwell system and infected at an MOI of 0.1 with parental SVN, R47, or mock inoculum. TEER was measured every 6 hours and media from the basolateral chamber was taken to measure viral titer every 6 hours. (B) Barrier integrity was measured as a function of transendothelial electrical resistance (TEER) at 6, 12, 18, or 24 hours post infection. Asterisks indicate statistically significant differences in TEER between mock and SVN infected (blue), mock and R47 infected (red), and SVN and R47 infected (black) hPSC-BMELCs at the indicated time points (two-way ANOVA and Šidák’s multiple comparisons): **, *p* < 0.01; ****, *p* < 0.0001. Non-significant differences are not shown. (C) Viral crossing was measured using the media harvested from the lower, basolateral chamber of the transwell system via plaque forming assay on BHK-21 cells. Asterisks indicate statistically significant differences in viral titer between SVN and R47 infected hPSC-BMELCs at the indicated time points (two-way ANOVA and Šidák’s multiple comparisons): ****, *p* < 0.0001. Non-significant differences are not shown. Data are representative of 2 independent experiments. Data is shown as mean ±SD.

### Neuroinvasive SINV is more efficient at hPSC-BMELC attachment, internalization, and fusion

Since BMEC infection is a critical correlate of neuroinvasive ability, we next wanted to identify the step in the virus infection cycle that is impacted by the neuroinvasive R47 specific mutations. Previous work demonstrated that the mutations in the E2 glycoprotein are critical for hPSC-BMELC infection^22^. Therefore, we wanted to investigate the role of E2 in hPSC-BMELC infection. We used single cycle pseudotyped vesicular stomatitis virus (VSV) particles to investigate the role of alphavirus entry in BMEC infection, similar to how these particles were used to investigate SARS-CoV-2 entry mechanisms^43^. Briefly, the alphavirus structural proteins will facilitate the VSV entry process, while the VSV components will replicate the genome containing a luciferase reporter gene. Importantly, the VSV genome does not encode any of its own structural genes and will therefore not produce any new particles. Thus, any difference in infectivity across VSV particles will be due to interactions of the alphavirus structural proteins with cellular factors at the entry steps. We cloned the SINV structural genes without the capsid gene and included an N-terminal myc tag to the E2 gene to generate VSV particles with SVN or R47 glycoproteins. We then ensured each VSV particle has similar amounts of E2 by measuring the amounts of myc-E2 in the virus stocks, therefore any differences in infection between the VSV particles will be due to genetic differences in the E2 glycoproteins between SVN and R47 (Figure S3A). Consistent with the role of E2 mutations in modulating alphavirus entry, the R47 pseudotyped VSV particles infect hPSC-BMELCs about 2-fold better than SVN pseudotyped particles, suggesting that SINV entry is the major determinant of BMEC infection (Figure 3A).

**Figure 3.**
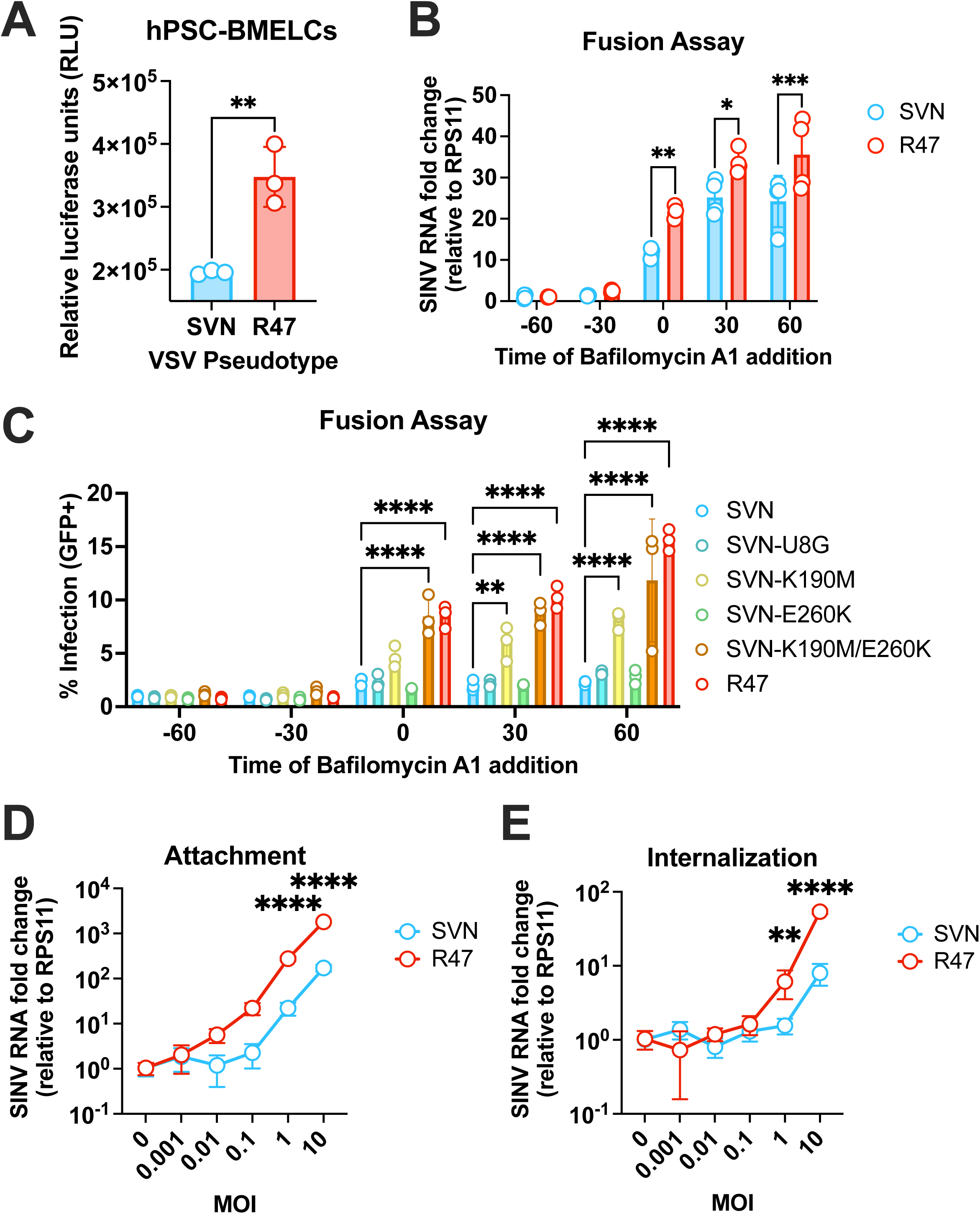
Neuroinvasive SINV is more efficient at hPSC-BMELC attachment, internalization, and fusion. (A) VSV particles pseudotyped with either SVN or R47 glycoproteins were used to infect hPSC-BMELCs at an MOI of 0.5. Cells were lysed 24 hours post infection and infection was measured via luciferase activity. Asterisks indicate statistically significant differences in viral titer between hPSC-BMELCs infected with either SVN- or R47- pseudotyped VSV particles (unpaired t-test): **, *p* < 0.01. Non-significant differences are not shown. (B and C), hPSC-BMELCs were treated with 25nM Bafilomycin A_1_ either 60 minutes before, 30 minutes before, at the same time, 30 minutes after, or 60 minutes after virus addition. (B) hPSC-BMELCs were infected with parental SVN or R47 at an MOI of 1 and harvested 12 hours post infection for RT-qPCR analysis. Asterisks indicate statistically significant differences in viral RNA fold change between SVN and R47 at each individual condition time point (two-way ANOVA and Šidák’s multiple comparisons): *, *p* < 0.05; **, *p* < 0.01; ***, *p* < 0.001. Non-significant differences are not shown. (C) hPSC-BMELCs were infected with GFP-expressing SVN, SVN point mutants, or R47 at an MOI of 0.1 and harvested 18 hours post infection for flow cytometry. Asterisks indicate statistically significant differences in infection with respect to SVN at each individual condition time point (two-way ANOVA and Šidák’s multiple comparisons): **, *p* < 0.01; ****, *p* < 0.0001. Non-significant differences are not shown. (D) SVN or R47 were incubated with hPSC-BMELCs at increasing MOIs at 4°C for 90 minutes. After several washes, bound virions were quantified as the ratio of the nsp1 gene to cellular RPS11A via RT-qPCR. Asterisks indicate statistically significant differences in attachment between SVN and R47 at each MOI (two-way ANOVA and Šidák’s multiple comparisons): ****, *p* < 0.0001. Non-significant differences are not shown. (E) Virions were allowed to internalize at 37°C for one hour in the presence of Bafilomycin A_1_. After removal of extracellular virions via proteinase K digestion, internalized virions were quantified as in (D). Asterisks indicate statistically significant differences in internalization between SVN and R47 at each MOI (two-way ANOVA and Šidák’s multiple comparisons): **, *p* < 0.01; ****, *p* < 0.0001. Non-significant differences are not shown. Data are representative of 2 independent experiments. Data is shown as mean ±SD.

For effective cell entry, alphaviruses must first 1) attach to the cell surface by binding to specific attachment factors and receptors, 2) be internalized into host endosomes, and 3) fuse with the endosomal membrane to release its genome. We hypothesize that R47 has a selective advantage over SVN during this viral entry process, which we assessed using a commonly applied entry assay^44,45^. Briefly, we treated hPSC-BMELCs with an inhibitor of alphavirus fusion, bafilomycin A_1_, at various time points prior to, during, and after virus addition. This assay allows us to identify differences in entry kinetics between SVN and R47 relative to the effects of bafilomycin A_1._ While bafilomycin A_1_ inhibits entry when added prior to virus addition, R47 RNA can be detected at high levels when bafilomycin A_1_ is added at the same time as virus, compared to SVN (Figure 3B). This is likely due to R47 having more efficient entry kinetics, which allows the virus to get in prior to the effects of bafilomycin A_1_, whereas SVN is not as efficient and is therefore fully inhibited by bafilomycin A_1_. This difference is emphasized as bafilomycin A_1_ is added at later time points, suggesting that R47 is more efficient in at least one of the entry steps leading up to and including fusion. While the mutations in E2 were shown to be important for hPSC-BMELC infection, we wanted to investigate whether and which of these mutations also facilitate more efficient entry. Therefore, we performed a similar assay with variants of SVN that have various individual and combined point mutations from R47. The first mutant, SVN-U8G displays no significant difference than SVN at any time point, suggesting this mutation at the 5’ UTR has no effect on entry and fusion (Figure 3C). Interestingly, SVN-E260K also has no significant difference compared to SVN at any time point, suggesting that the residue at position 260 alone is not sufficient to increase entry efficiency (Figure 3C). By contrast, SVN-K190M enters hPSC-BMELCs significantly more efficiently starting when bafilomycin A_1_ is added 30 minutes after virus addition, suggesting that the K190M mutation is critical for the entry advantage over SVN; however, the advantage is delayed and not to the same degree as R47 (Figure 3C). Interestingly, the E260K mutation confers an additional advantage in the background of the K190M mutation and together match the entry efficiency as R47 (Figure 3C). These data are consistent with *in vivo* experiments demonstrating that the K190M change is the critical mutation that drastically decreases the intraperitoneal LD50 compared to SVN, but both K190M and E260K mutations are required in the background of the U8G mutation for reaching the same intraperitoneal LD50 as R47^7^. Together, this work suggests that the 190 position is critical for BMEC entry and neuroinvasion while the 260 position increases the replicative fitness of the virus in BMECs, leading to enhanced neuroinvasion.

Our work thus far demonstrates that R47 E2 confers an advantage at viral entry, which is critical for BMEC infection. However, to thoroughly understand the critical interactions we must determine the entry step that is initially exploited by R47, which include cell attachment and internalization. For attachment, we allowed varying concentrations of virus particles to bind to hPSC-BMELCs for one hour and measured viral RNA immediately afterward. Interestingly, R47 attaches more efficiently than SVN starting at the MOI of 0.01, however this difference does not become statistically significant until MOI of 1 (Figure 3E). At higher concentrations, it takes ten times as many SVN particles to get the same level of attachment as R47 (Figure 3E). For internalization, we allowed SVN or R47 particles to become internalized within hPSC-BMELCs in the presence of bafilomycin A_1_ to prevent fusion. Additionally, we treated hPSC-BMELCs with proteinase K to remove plasma membrane bound, uninternalized virions, to allow us to measure only changes in internalization. Similarly to the binding assay, we measured viral RNA as a proxy for internalized virions. We found that SVN and R47 internalize similarly at low MOIs but begin to significantly differ at MOIs of 1 and 10 (Figure 3D). These data demonstrate that both cell attachment and internalization are highly critical for the efficient infection of BMECs. Since the largest difference between SVN and R47 is during hPSC-BMELC attachment, we decided to first investigate a well-known attachment factor: heparan sulfate. We first pretreated hPSC-BMELCs with heparanases which remove heparan sulfate from the cell surface prior to infection with SVN and R47. To our surprise, removal of heparan sulfate is not sufficient to explain the differential infection between SVN and R47 as there is still a >2-fold difference between the strains (Figure S3B). Therefore, these data suggest that other host entry factors may be exploited for the efficient infection of BMECs, such as receptors, which are highly involved throughout the entry process.

### No known receptor is uniquely engaged by R47

We have demonstrated that neuroinvasive SINV must efficiently attach and enter BMECs for high levels of BMEC infection; however, the host factors exploited during this mechanism remain unknown. We have also shown that the differential infection phenotype is not dependent on heparan sulfate, therefore we investigated the role of cell surface protein receptors. SINV is known to interact with a handful of previously identified receptors including SLC11A2, avian MXRA8, VLDLR, and LRP8^32,35,37^; however, none of these have been studied in the context of neuroinvasion, and specifically, infection of the BBB. To narrow down our list of candidates, we first performed RNA-sequencing followed by differential transcriptomic analysis on hPSC-BMELCs and HUVECs under basal conditions. We then performed quality control steps and found two of the HUVEC samples to be different from the rest of the replicates through sample-to-sample heatmap analysis (Figure S4A); however, when the samples were plotted via principal component analysis (PCA) the difference between those samples and the rest only accounts for 2% of the total variance across the experiment (Figure S4B, refer to the y-axis PC2), and therefore the samples were kept in the analysis. Since HUVECs were efficiently infected by both SVN and R47, but hPSC-BMELCs were only efficiently infected by R47, we hypothesized that the host factor exploited by R47 must be uniquely engaged by R47 and abundantly expressed in hPSC-BMELCs but not HUVECs.

From our analysis, we identified >8000 differentially expressed genes (DEGs) between hPSC-BMELCs and HUVECs. All the known alphavirus receptors are among the significant DEGs, except for MXRA8, which was excluded due to insufficient read coverage. Among the DEGs encoding alphavirus receptors, only VLDLR and SLC11A2 fit our criteria as being most abundant in hPSC-BMELCs but not HUVECs (Figure 4A). To assess whether SVN or R47 could engage either of these receptors, we first performed receptor blockade assays in HEK293T cells. Although HEK293T cells are not representative of the BBB, they are commonly used in alphavirus studies due to their permissiveness to infection and are known to express several known alphavirus receptors. This simplified system allowed us to assess whether SVN and R47 could engage with the candidate receptors in a more controlled context. Blockade of VLDLR did not significantly affect either SVN or R47 infection, indicating that this receptor is not critical for viral entry in these cells. In contrast, blockade of SLC11A2 resulted in an 8-fold reduction in infection by both SVN and R47, suggesting that SLC11A2 is involved in viral entry (Figure 4B). However, because SLC11A2 was engaged by both viruses, it does not meet our criteria as a receptor uniquely exploited by R47 for neuroinvasion.

**Figure 4.**
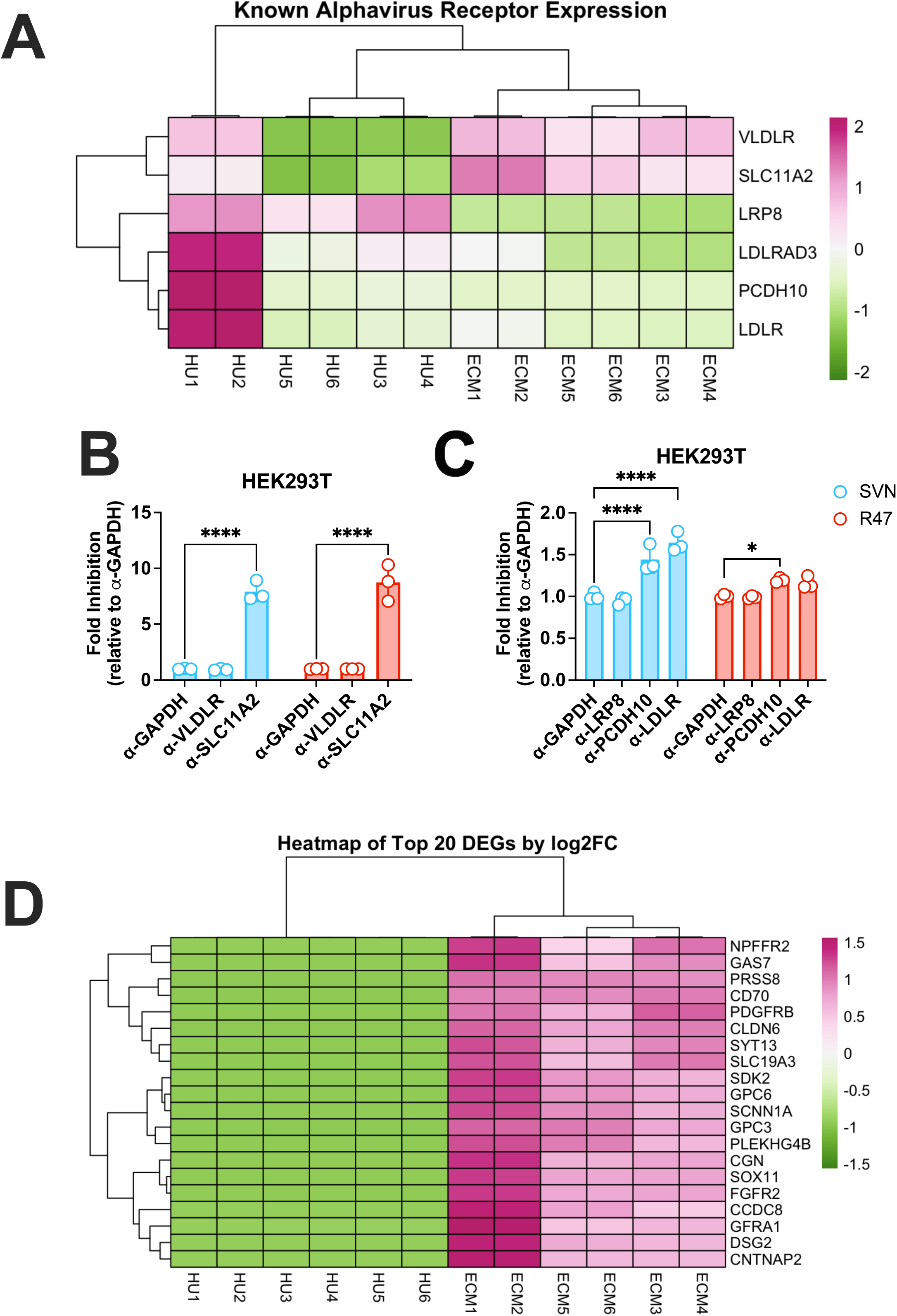
No known receptor is uniquely engaged by R47. hPSC-BMELCs and HUVECs were independently cultured, and total RNA was isolated for RNA-seq and data analysis using DESeq2. (A) Heatmap by log2FC of previously identified alphavirus receptors that were significantly enriched in both hPSC-BMELCs and HUVECs. (B) HEK293T cells were pretreated with antibodies targeting either GAPDH, VLDLR, or SLC11A2 an hour before infection with GFP-expressing SINV strains of SVN or R47 at an MOI of 1. Cells were harvested 18 hours post infection and analyzed via flow cytometry. Asterisks indicate statistically significant differences in fold-inhibition with respect to the control antibody (two-way ANOVA and Šidák’s multiple comparisons): ****, *p* < 0.0001. Non-significant differences are not shown. (C) HEK293T cells were pretreated with antibodies targeting either GAPDH, LRP8, PCDH10, or LDLR an hour before infection with GFP-expressing SINV strains of SVN or R47 at an MOI of 1. Cells were harvested 18 hours post infection and analyzed via flow cytometry. Asterisks indicate statistically significant differences in fold-inhibition with respect to the control antibody (two-way ANOVA and Šidák’s multiple comparisons): *, *p* < 0.05; ****, *p* < 0.0001. Non-significant differences are not shown. Data are representative of 2 independent experiments. Data is shown as mean ±SD. (D) Heatmap of the top 20 differentially expressed genes in hPSC-BMELCs with a log2FC>0 compared to HUVECs. Hits that overlap with transcriptomic datasets are indicated by asterisk.

While the remaining known receptors are not highly expressed in hPSC-BMELCs compared to HUVECs, we decided to test whether any of these remaining receptors are uniquely engaged by R47 since it is possible that even low expression of these receptors on BMECs may facilitate a differential infection phenotype. Specifically, we focused on receptors that have been shown to be engaged by SINV and the SINV-related Western equine encephalitis virus, including LRP8, PCDH10, and LDLR. Interestingly, blockade of LDLR and PCDH10 results in modest yet significant inhibition of SVN replication in HEK293T cells, while R47 was only inhibited by PCDH10 blockade but to a lesser degree (Figure 4C). Together these data suggest that both SVN and R47 can use SLC11A2 and PCDH10 to a lesser extent, while only SVN can also use LDLR as a receptor for infecting HEK293T cells. Ultimately, none of the previously identified alphavirus receptors are likely candidates that may explain the efficient infection of BMECs by R47 since none of these receptors are uniquely engaged by R47 and highly upregulated in hPSC-BMELCs.

We hypothesized that the transcriptomic data may yield some insight towards identifying a novel receptor candidate that is specific to hPSC-BMELCs and uniquely engaged by R47. We refined our initial list of DEGs by performing gene ontology (GO) analysis by cellular component, and selected DEGs that have GO terms associated with the cell surface, resulting in over 3000 DEGs. We filtered even further and selected only the DEGs that are more abundant in hPSC-BMELCs than HUVECs, which yields just over 1000 DEGs. Among the top 20 candidates are transcripts encoding tight junction proteins important for maintaining the BBB like claudin 6 (CLDN6) and cingulin (CGN), which is expected considering hPSC-BMELCs are an endothelial cell model for cells that form tight barriers compared to HUVECs. Additionally, we found other transcripts encoding proteins related to known alphavirus receptors like desmoglein 2 (DSG2; a cadherin protein related to the PCDH10 receptor), and solute carrier family 19 member 3 (SLC19A3; a transporter protein in the same family as SLC11A2) (Figure 4D). While these hits are promising, there are still over 1000 possible candidates; therefore, complementary datasets must be generated to further refine this list of novel receptor candidates.

### Cell surface proteomic analysis reveals 4 novel alphavirus receptor candidates

To refine our list of candidate receptors and corroborate these findings on the protein level, we performed cell surface proteomics on hPSC-BMELCs, HUVECs, and HCMEC/D3 cells. Since HCMEC/D3 cells are not susceptible to SVN or R47 (Figure S2D), we included these cells in the analysis as a negative control. Among these three endothelial cell types, the host entry receptor is likely unique to hPSC-BMELCs and will be identified among the significantly upregulated proteins in both hPSC-BMELCs vs HUVECs comparison and hPSC-BMELCs vs HCMEC/D3 cells comparison. Additionally, while all three cell types are expected to share endothelial cell features, hPSC-BMELCs and HCMEC/D3 cells are of the brain origin and therefore should have the greatest overlap in endothelial cell proteomes. We labeled cell surface proteins with biotin in the three endothelial cell types, enriched for biotinylated proteins using neutravidin beads, and subjected the precipitates to proteolytic digestion and mass spectrometry analysis. Indeed, upon comparing our cell surface proteomes via PCA, we found that hPSC-BMELCs cluster closely with HCMEC/D3 cells compared to HUVECs along PC1, which explains the greatest variance between samples (Figure S5A, refer to x-axis). Moreover, the hPSC-BMELCs are also slightly distinct from the HCMEC/D3 cells, yet still endothelial cell-like since the hPSC-BMELCs cluster among HUVECs while being distinct from HCMEC/D3 cells along PC2 (Figure S5A, refer to y-axis). Our enrichment was successful as demonstrated by a 5-fold increase in unique peptides and a 3-fold increase in unique proteins in the biotinylated samples compared to the non-biotinylated samples (Figure 5A and 5B). Moreover, when the samples were plotted via PCA, the biotinylated samples cluster farther away from the non-biotinylated samples and explains almost 80% of the variance, which is indicative of efficient enrichment (Figure S5B). Together these data give us high confidence that our biotinylation reactions are successful and that we enriched for cell surface proteins in each of our endothelial cell types.

**Figure 5.**
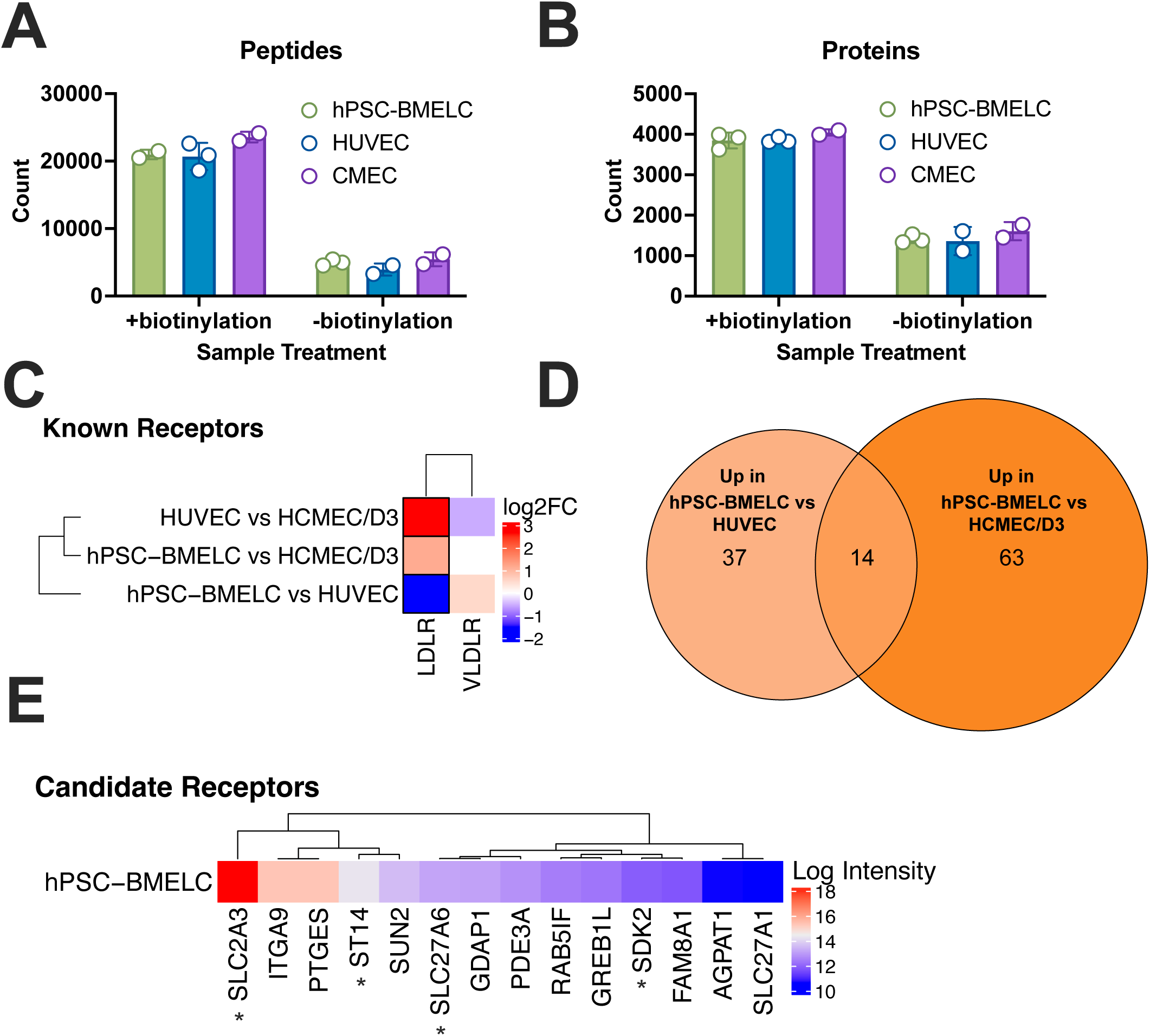
Cell surface proteomics analysis reveals 4 novel alphavirus receptor candidates. hPSC-BMELCs, HUVECs, and HCMEC/D3 cell surfaces were biotinylated, and surface proteins were harvested for proteomic analysis. (A-B) Column graph representing the total number of unique peptides (A) or unique proteins (B) enriched in either biotinylated or non-biotinylated samples. (C) Heatmap of previously identified alphavirus receptors that were detected in any of the three endothelial cell types. (D) Venn diagram of proteins with a transmembrane domain that are most abundant in hPSC-BMELCs compared to HUVECs or HCMEC/D3. (E) Heatmap showing relative protein intensity in hPSC-BMELCs of the 14 proteins that were found to be uniquely abundant in hPSC-BMELCs compared to HUVECs and HCMEC/D3 cells.

We took our enriched protein list and refined it by filtering out the proteins that do not have gene ontology terms associated with a transmembrane domain. Similar to our transcriptomic analysis, we first wanted to interrogate whether any known alphavirus receptors were detected in hPSC-BMELCs and fit our criteria of being most abundant in hPSC-BMELCs compared to the other endothelial cell types. Among the enriched proteins, LDLR and VLDLR are the only known alphavirus receptors that were detected among our samples (Figure 5C). However, neither fit our expected pattern, and instead both LDLR and VLDLR are mostly enriched in the other endothelial cell types compared to hPSC-BMELCs. Therefore, these receptors may be engaged during infection of HUVECs but not hPSC-BMELCs.

Since we hypothesize that the host entry factor exploited by R47 should be highly upregulated in hPSC-BMELCs, we looked into proteins that are more abundant in hPSC-BMELCs compared to HUVECs and HCMEC/D3 cells. In total, 51 proteins are significantly enriched in hPSC-BMELCs over HUVECs and have gene ontology terms associated with a transmembrane domain, and 77 proteins are significantly enriched in hPSC-BMELCs over HCMEC/D3 cells that have gene ontology terms associated with a transmembrane domain (Figure 5D). Among these proteins, 14 overlap in our comparisons, suggesting that the host factor exploited by R47 is likely within this list. When we plotted the relative protein intensity from hPSC-BMELCs of these 14 shared proteins, we found solute carrier family 2 member 3 (SLC2A3, also named GLUT3) to be the most highly expressed and unique to hPSC-BMELCs (Figure 5E). To refine this list of candidates further, we overlayed our transcriptomic analysis and our proteomic analysis, which results in 4 novel receptor candidates: SLC2A3, solute carrier family 27 member 6 (SLC27A6, also named FATP6), sidekick 2 (SDK2), and transmembrane serine protease matriptase (ST14) (Figure 5E, asterisks). Members of the SLC family are among the most promising candidates as they are structurally related to SLC11A2, which also has the strongest effect on SVN and R47 infection (shown in Figure 4B). Importantly, SLC2A3/GLUT3 is uniquely found on neuronal cells and cells of the BBB, while SLC27A6/FATP6 is primarily found in the heart^46,47^. Together, our data point to SLC2A3 as the most likely candidate receptor for R47 exploitation since SINV engages a highly related receptor, SLC11A2, and SLC2A3 is uniquely and highly expressed in cells of the BBB including BMECs.

### BMEC infection correlates with neuroinvasive ability of diverse Old World alphaviruses

We sought to determine whether the viral determinants of neuroinvasion are shared among other Old World alphaviruses. We first infected hPSC-BMELCs with several Old World alphaviruses, including ONNV, RRV, and CHIKV in addition to SVN and R47. We found that none of the additional Old World alphaviruses infect hPSC-BMELCs beyond 1% (Figure 6A). However, among these alphaviruses, only CHIKV is known to cause severe complications of the central nervous system via potential direct neuroinvasion, as evidenced by the CHIKV RNA found in the cerebrospinal fluid of fatal human cases. Furthermore, *in vivo* mouse studies have demonstrated that the CHIKV 181/clone 25 vaccine strain is non-neuroinvasive while the virulent La Reunion (LR) strain is neuroinvasive. Therefore, we investigated whether there would be a differential infection phenotype of CHIKV strains in cells of the BBB similar to SVN and R47. Indeed, we found that the virulent CHIKV LR strain efficiently infects hPSC-BMELCs 6-fold more than the attenuated 181/clone 25 vaccine strain. Moreover, the CHIKV LR strain significantly infects hPSC-BMELCs (40%) to similar levels as R47 (55%) (Figure 6A and 6B). Additionally, pericytes are infected 2-fold more by the LR strain than the 181/clone 25 vaccine strain. Importantly, astrocytes are efficiently infected by both CHIKV strains, suggesting the previous phenotypes are not a result of broad attenuation (Figure 6B). Together these data support the hypothesis that neuroinvasive Old World alphaviruses have the ability to efficiently infect all cell types of the BBB.

**Figure 6.**
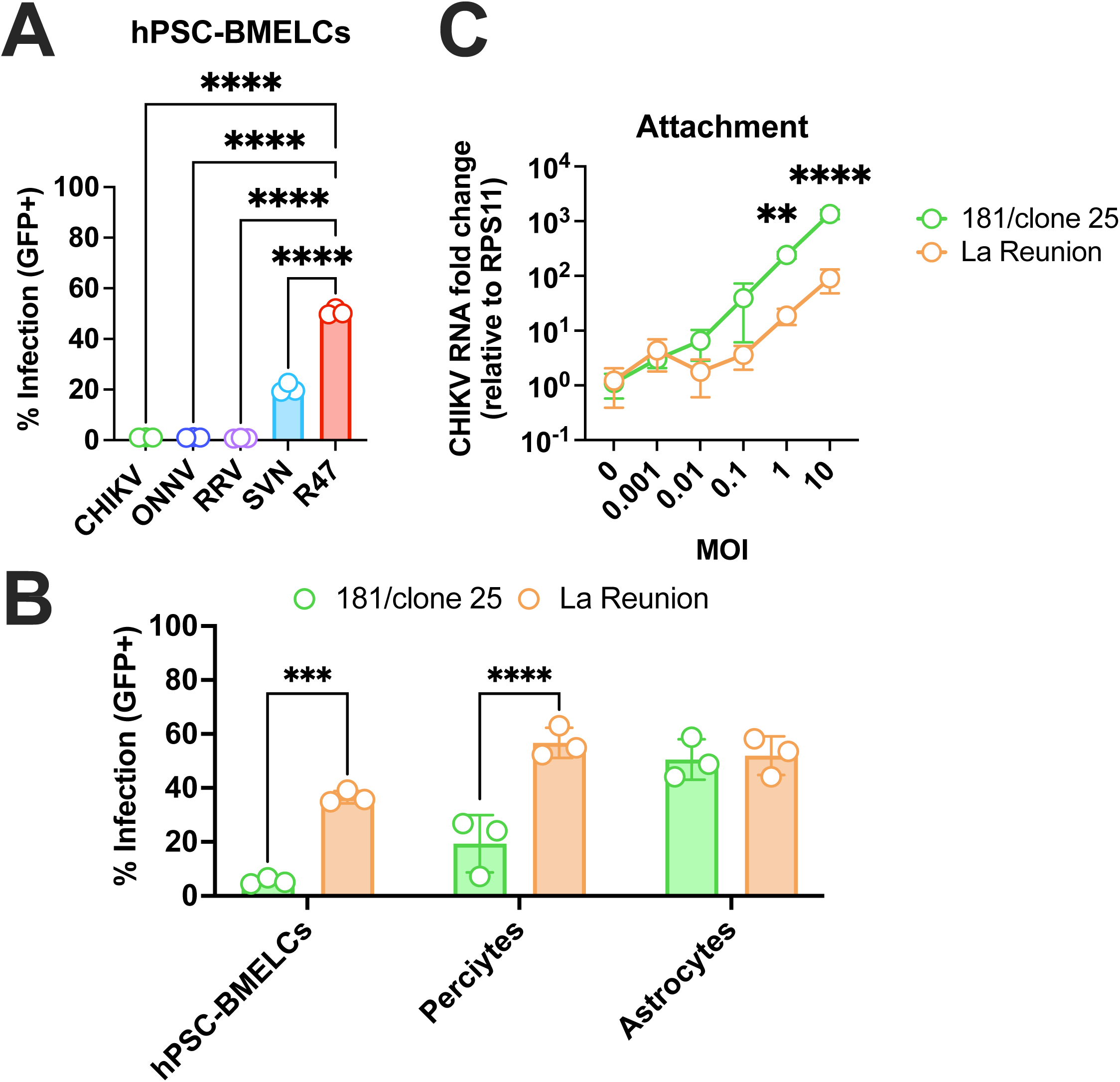
BMEC infection correlates with neuroinvasive ability of diverse Old World alphaviruses. (A) hPSC-BMELCs were infected at an MOI of 0.1 with GFP-expressing chikungunya virus vaccine strain 181/clone 25 (CHIKV 181/clone 25), o’nyong’nyong virus (ONNV), Ross River virus (RRV), SINV strain SVN, or SINV strain R47. Cells were harvested 24 hours post infection and were analyzed via flow cytometry. Asterisks indicate statistically significant differences in infection with respect to R47 (ordinary one-way ANOVA and Dunnett’s multiple comparisons): ****, *p* < 0.0001. Non-significant differences are not shown. (B) hPSC-BMELCs, primary pericytes, or primary astrocytes were infected with GFP-expressing CHIKV vaccine strain 181/clone 25 or CHIKV virulent strain La Reunion (LR) at an MOI of 0.1. Cells were harvested 40 hours post infection and analyzed via flow cytometry. Asterisks indicate statistically significant differences in infection between 181/clone 25 and the LR strain in the indicated cell type (two-way ANOVA and Šidák’s multiple comparisons): ***, *p* < 0.001; ****, *p* < 0.0001. Non-significant differences are not shown. (C) CHIKV 181/clone 25 or LR GFP-expressing viruses were incubated with hPSC-BMELCs at increasing MOIs at 4°C for 90 minutes. After several washes, bound virions were quantified as the ratio of the nsp1 gene to cellular RPS11A via RT-qPCR. Asterisks indicate statistically significant differences in attachment between 181/clone 25 and LR at each MOI (two-way ANOVA and Šidák’s multiple comparisons): **, *p* < 0.01; ****, *p* < 0.0001. Non-significant differences are not shown. Data are representative of 2 independent experiments. Data is shown as mean ±SD.

Lastly, we wanted to investigate whether BMEC attachment is the critical interaction necessary for CHIKV neuroinvasion as it is for SINV. We performed the same binding assay as before but with the CHIKV 181/clone 25 vaccine strain and LR strain. Surprisingly, the CHIKV LR strain binds to hPSC-BMELCs almost 10 times less than the CHIKV 181/clone 25 vaccine strain (Figure 6C). This suggests that, while neuroinvasive Old World alphaviruses have gained the ability to infect cells of the BBB for neuroinvasion, they have evolved distinct mechanisms for infection that may involve multiple stages in the viral life cycle. Together, these data identify BBB infection as critical for subsequent neuroinvasion for SINV and the more clinically relevant CHIKV. This can potentially provide a broad-spectrum target for blocking Old World alphavirus entry into and infection of the CNS.

## DISCUSSION

Our study is the first to our knowledge to establish a correlation between BBB infection, neuroinvasion, and alphavirus pathogenesis. By using hPSC-BMELCs to investigate SINV-BBB molecular interactions, we show that neuroinvasive E2 residues confer the ability to efficiently infect BMECs and pericytes. This phenotype is driven by just two residues on E2, which critically enhance BMEC attachment and entry. Moreover, we show that none of the previously identified alphavirus receptors are differentially engaged by SVN and R47, and we propose SLC2A3 as a novel receptor candidate that is exploited by R47 for neuroinvasion. Strikingly, enhanced BBB infection was also shown to strongly correlate with the neuroinvasive ability of the virulent La Reunion strain of CHIKV. However efficient infection is not dependent on enhanced attachment, suggesting SINV and CHIKV have evolved distinct mechanisms to achieve efficient BBB infection. The data ultimately demonstrate that infection of the blood-brain barrier may be essential for pathogenesis.

Stem cell-derived models of viral infection are critical for driving our understanding of viral pathogenesis forward by enabling studies in a more physiologically relevant context^48–51^. For instance, hPSC-BMELCs have been pivotal in investigating flavivirus neuroinvasion, revealing key host factors that restrict certain flaviviruses and prevent their entry into the CNS^22^. hPSC-BMELCs address a longstanding challenge in BBB modeling, as traditional approaches have significant limitations. Primary BMECs are difficult to isolate and culture beyond a few passages, and immortalized cell lines, such as HCMEC/D3, are easier to culture but lack the robust barrier integrity required for transwell systems. By contrast, hPSC-BMELCs are easily generated in large quantities, offer physiologically relevant barrier properties, and can be genetically manipulated. Despite some differentiation protocols yielding BMEC-like cells that also express some non-endothelial markers^18^, ongoing advancements—such as the incorporation of retinoic acid to enhance barrier integrity, shear force to better mimic vascular conditions, and the introduction of specific transcription factors—continue to refine these models^16,17,52–54^. This adaptability underscores the unique advantage of stem cell-derived systems, which can evolve to address experimental needs. To complement our work, future studies aim to incorporate viral infection in more physiological BBB models, such as introducing flow to investigate how efficient BMEC attachment changes in circulating fluid and natural vascularization to better understand dissemination from the BBB to the brain parenchyma.

Surprisingly, our model revealed a neuroinvasion mechanism that has not been observed *in vivo*. Previous work on VEEV and WEEV neuroinvasion suggest that cells of the murine BBB are not infected *in vivo*^12^. Instead, VEEV transverses BMECs via caveolin-mediated endocytosis, implying that direct BBB infection is not a conserved neuroinvasion strategy among encephalitic alphaviruses. While these studies showed that all murine BBB cell types and human BMECs are susceptible and permissive to VEEV and WEEV infection *in vitro*, no evidence of BMEC or pericyte infection was found *in vivo* in the presence of type I IFN signaling. This discrepancy could be a result of the time points chosen, as infection may occur transiently or at earlier stages of disease. Additionally, mechanisms of neuroinvasion in adolescent mice may differ from mice representative of populations most affected by severe disease, such as young children or older adults. An alternative explanation is that New World alphaviruses have evolved distinct mechanisms for neuroinvasion compared to Old World alphaviruses, underscoring the complexity and diversity of alphavirus pathogenesis. To address this, future work may investigate the ability to infect BMECs across neuroinvasive and non-neuroinvasive strains of New World alphaviruses, such as the attenuated VEEV TC83 and the virulent VEEV TrD.

Mechanisms of SINV and CHIKV neuroinvasion have remained elusive, despite their critical importance in the context of CHIKV-associated fatal infections^4^. Several *in vivo* models, including zebrafish, mice, and human cohorts, have been employed to address this question. Zebrafish studies have demonstrated that neither SINV nor CHIKV rely on macrophages to traverse the BBB, while murine models reveal that SINV crosses the BBB before any observable barrier disruption^13,14^. Human cohort studies suggest that CHIKV crosses the BBB through leakage and immune cell carriage, findings that contrast with those from zebrafish models and our *in vitro* model^4^. Notably, while BMEC infection was not observed in human samples, this does not entirely exclude the possibility of BMECs serving as a route for neuroinvasion. It is technically difficult to determine whether BMECs are infected in human cases since by the time the individual is dead the infected cells might be killed. These discrepancies underscore the limitations of current models, as *in vivo* studies often capture data during late stages of disease or post-mortem, overlooking the immune-mediated nature of encephalitic disease^55–57^. Our stem cell derived model provides a powerful platform for dissecting molecular mechanisms to complement murine studies that are critical for understanding complex cell-cell interactions in the context of the whole organism. Together, these findings highlight the importance of using multiple models for investigating neuroinvasion. Therefore, to complement our study, future studies looking at the role of BMEC infection *in vivo* will be highly informative.

Our study is among the first to investigate the role of host entry factors in relevant human cell types that are critical for alphavirus pathogenesis. Previous studies identified several low-density lipoprotein receptor (LDLR) proteins as essential receptors for several encephalitic alphaviruses including VEEV, WEEV, EEEV, and SFV^32–34^. For SINV, the LDLR-related proteins LRP8 and VLDLR have been implicated as receptors in mouse neuroblastoma N2A cells, although this appears to be context-dependent. For instance, while SINV structural proteins can bind LRP8 and VLDLR, overexpression of these receptors in non-susceptible cell lines only modestly increases infection rates to approximately 20%. Furthermore, blocking these receptors on susceptible cells does not significantly reduce SINV infection, in contrast to SFV, for which infection is entirely abrogated. Other groups have shown that several Old World alphaviruses, including CHIKV, ONNV, and RRV, use MXRA8 as a receptor on mouse embryonic fibroblasts and 3T3 cells, which is critical for mouse pathogenesis^31^. Interestingly, SINV, despite being an Old World alphavirus, does not rely on mammalian MXRA8, but rather avian MXRA8 since wild birds are the natural reservoir for SINV^35,58^. The most compelling work that identified a receptor for SINV determined that SLC11A2 is a cell entry receptor for SINV in mosquito cells and primary murine fibroblasts^37^; however, it is unclear how cell type specific expression of SLC11A2 contributes to pathogenesis. Our data highlights SLC11A2 as an entry factor that can be used by both SVN and R47, likely for the cell types they both infect equally, such as HUVECs. When investigating cell receptors that facilitate neuroinvasion, our work suggests that SLC2A3 could be a receptor engaged by neuroinvasive variants of SINV. Importantly, this receptor is abundant in the brain and specifically on BMECs^46^. This finding is especially striking since SLC2A3 is within the same family as SLC11A2 and shares some structural homology, making SLC2A3 a promising candidate for further validation and future therapeutic development. Altogether, these data suggest that SINV uses multiple receptors efficiently while other alphaviruses prefer a few receptors. This interesting phenotype raises important questions about how SINV receptor usage influences its overall pathogenesis. Future studies must focus on validating the role of SLC2A3 in BMEC infection to uncover how these potential interactions contribute to SINV’s unique biology and pathogenesis, and our ongoing work aims to address these gaps.

Cell tropism plays a critical role in shaping pathogenesis and disease outcomes, particularly in fatal cases of CHIKV. For instance, our work demonstrates that neuroinvasive alphaviruses exploit host entry factors, driving BMEC tropism, and leading to neuropathogenesis. Moreover, this work is underscored by previous findings showing CHIKV RNA in the cerebrospinal fluid of all fatal CHIKV cases, suggesting neuroinvasion is a key determinant of lethality^4^. These insights emphasize the importance of targeting BBB-specific receptors to prevent alphavirus neuroinvasion and its associated fatal outcomes. Therapeutic strategies include receptor-targeting antibodies or soluble receptor therapeutics that block viral entry into barrier cells, thereby preventing neuroinvasion.

Our findings also open new avenues for vaccine development. The determinants of attenuation for vaccine strains of CHIKV and VEEV were previously mapped to mutations in the E2 glycoprotein, believed to attenuate the virus through increased affinity for heparin sulfate^29,59^. However, the functional consequences of these mutations on tropism are not well understood. We reveal that attenuating mutations in CHIKV result in a loss of BMEC and pericyte tropism, directly linking neuroinvasion and pathogenesis. Follow-up studies should aim to validate these findings across additional alphaviruses, demonstrating that barrier tropism is a primary determinant of pathogenesis *in vivo*. This suggests that future attenuated vaccines for other alphaviruses could be designed to specifically disrupt barrier tropism, thereby reducing virulence.

This study has found that BBB infection correlates with the neuroinvasive and pathogenic phenotypes of several alphaviruses. Notably, we show here that loss of BBB infection may provide a molecular mechanism for viral attenuation. Ultimately, therapeutic approaches based on our work will not only help treat ongoing cases of CHIKV neuroinvasion, but builds a potential fast-track for vaccine design against an increasingly plausible alphavirus pandemic.

## LIMITATIONS OF THE STUDY

We acknowledge three primary limitations of our study with regards to our hPSC-derived model, applications *in vivo*, and the systematic investigation of CHIKV neuroinvasion. First, the hPSC-BMELCs used in this study are derived using recent and advanced differentiation protocols; however, these endothelial-like cells are known to also express some epithelial cell markers^18^. Future experiments should increase the model’s accuracy by including co-cultures of other BBB cells and potentially including vascularized 3D cultures. Second is the lack of *in vivo* work that further corroborates the importance of BMEC infection in SINV and CHIKV neuropathogenesis. We have shown that infection of the BBB strongly correlates with neuroinvasive capacity, but future work *in vivo* should determine whether the murine BBB is efficiently infected via immunostaining. Additionally, future work should investigate the role of SLC2A3 on R47 neuroinvasion in mice. Finally, we acknowledge the limitations of our system to investigate CHIKV neuroinvasion. In this study, we used two strains of CHIKV with opposite *in vivo* neuroinvasiveness; however, using just two strains limits our ability to pinpoint residues critical for infection of the BBB. Instead, future work will apply evolutionary approaches to systematically identify residues critical for CHIKV neuroinvasion. The limitations of our study present exciting opportunities for future research that will facilitate the next step in the development of novel therapeutic strategies to mitigate alphavirus induced neuropathies.

## ACKNOWLEDGEMENTS

We thank Faith St. Amant, Sangeetha Ramachandran, and Dr. Zhenlan Yao, for their support, critical discussions, expertise and advice that facilitated the successful construction of myc-tagged alphavirus glycoproteins. We thank Dr. LeAnn Nguyen, Dr. Serina Huang, Erin Kim, and Martin Ruvalcaba for their support, expertise, and critical discussions that helped push this work forward. We thank Dr. Michael Letko (Washington State University) and Dr. Vincent Munster (National Institute for Allergy and Infectious Diseases) for providing the protocols and VSV-ΔG seed particles necessary for generating SINV pseudoviruses. We thank Dr. Charles M. Rice at the Rockefeller University, Dr. Stephen Higgs at Kansas State University, Dr. Scott Weaver at the University of Texas Medical Branch at Galveston, and Dr. Mark Heise at the University of North Carolina at Chapel Hill for providing the alphavirus constructs used in this study. We thank the QCBio Collaboratory at the Institute for Quantitative and Computational Biosciences for their services. We thank the James B Pendleton Charitable Trust and the McCarthy Family Foundation for providing the UCLA AIDS Institute with RT-qPCR and flow cytometry platforms. This work was funded in part by the UCLA Broad Stem Cell Research Center (BSCRC) Research Award (to MMHL), the UCLA BSCRC Rose Hills Foundation Innovation Award (to MMHL), a 2023 UCLA DGSOM W. M. Keck Foundation Junior Faculty Award (to MMHL), the National Institute of Allergy and Infectious Diseases of the National Institutes of Health (NIH) under Award Number F31AI179235 (to PAA), and the Ruth L. Kirschstein National Research Service Award under Award Number AI007323 (to PAA). The content is solely the responsibility of the authors and does not necessarily represent the official views of the NIH.

## DECLARATION OF INTERESTS

The authors declare no competing interests.

## STAR METHODS

### RESOURCE AVAILABILITY

#### Lead contact

Requests for resources, reagents, and further information should be directed to the lead contact, Melody M.H. Li.

#### Materials availability

Requests for resources and reagents, including viruses and cells, should be made to the lead contact author. All reagents and resources will be made available after completion of a Materials Transfer Agreement (MTA).

#### Data and code availability

All data from this study are available within the paper and from the lead contact author upon request. This paper does not include original code. RNA sequencing data have been deposited in the Gene Expression Omnibus (GEO) database. The mass spectrometry proteomics data have been deposited to the ProteomeXchange Consortium via the PRIDE^60^ partner repository with the dataset identifier PXD059899. The RNA-Sequencing data discussed in this publication have been deposited in NCBI’s Gene Expression Omnibus^61^ and are accessible through GEO Series accession number GSE287658.

### EXPERIMENTAL MODEL DETAILS

#### Cells. Human Pluripotent Stem Cells

Human embryonic stem cell line H9 (WA09; female) and human induced pluripotent stem cell line IMR90-4 (WiCell; female) were thawed in flasks coated with 0.2mg/mL Matrigel™ in DMEM/Ham’s F-12 (Gibco, 11320033). Stem cells were cultured in mTeSR™1 (STEMCELL Technologies, 85857) or mTeSR™ plus (STEMCELL Technologies, 100-0276). For all the experiments in this study, H9 cells were used between passages 48 – 60, and IMR90-4 cells were used between passages 38-42.

#### Primary Human Cells

Primary human umbilical vein endothelial cell (HUVECs; ATCC, PCS-100-010) were cultured in complete Endothelial Cell Medium (Millipore Sigma, 211-500) at 37°C and 5% CO_2_. Primary cortical human astrocytes (ScienCell, 1800) were cultured in poly-D-lysine (2 μg/cm^2^) coated culture vessels using supplemented astrocyte media (ScienCell, 1801) according to the provider’s instructions at 37°C and 5% CO_2_. Primary brain vascular pericytes (ScienCell, 1200) were cultured in Poly-D-lysine (2 μg/cm^2^) coated culture vessels using supplemented pericyte media (ScienCell, 1201) according to the provider’s instructions at 37°C and 5% CO_2_.

#### Immortalized Cell Lines

HCMEC/D3 cells were cultured on flasks coated with type I rat tail collagen using Endo-Gro-MV medium supplemented with 1ng/mL human basic fibroblast growth factor. Vero and HEK-293T cells were cultured in Dulbecco’s modified Eagle’s medium (DMEM) supplemented with 10% FBS at 37°C and 5% CO_2_. Baby hamster kidney (BHK-21) cells were cultured in MEM supplemented with 7.5% FBS at 37°C and 5% CO_2_.

##### Viruses

SINV SVN, R47, SVN-expressing GFP, and R47-expressing GFP (gift from Dr, Charles M. Rice, Rockefeller University)^7,13,22^, RRV expressing EGFP (gift from Dr. Mark Heise, University of North Carolina)^62^, ONNV expressing EGFP and CHIKV La Reunion strain expressing GFP (gift from Dr. Stephen Higgs, Kansas State University)^63,64^, CHIKV vaccine strain 181/clone 25 expressing EGFP (gift from Scott Weaver, The University of Texas Medical Branch at Galveston)^63^, have been previously described^65–67^. All alphavirus stocks were generated and viral titers were calculated using BHK-21 cells^68^. The amount of virus used for each experiment was determined by the multiplicity of infection (MOI), cell number, and virus titer.

### METHOD DETAILS

#### Generation of hPSC-BMELCs from Human Pluripotent Stem Cells

Stem cells were differentiated into hPSC-BMELCs according to previous publications (Lippmann et al., 2012; Lippmann et al., 2014; Neal et al., 2019). Briefly, stem cells were thawed on Matrigel coated flasks and grown for a minimum of 4 days in mTeSR1. After the fourth day, mTeSR1 was replaced by growth media (20% KnockOut Serum Replacer, 1mM L-glutamine, 1X NEAA, 0.1mM β-mercaptoethanol, and 100ng/mL basic fibroblast growth factor in 1:1 DMEM/F-12) to initiate differentiation. After 5-7 days, the growth media was changed to endothelial cell (EC) media (Gibco, 11111044) supplemented with 1X B-27 supplement, 20 ng/mL human basic fibroblast growth factor and 10μM of retinoic acid (RA). After 2 days, the cells were dissociated with StemPro Accutase (Gibco, A1110501) for 30-45 minutes at 37°C, followed by cell scraping. Cells were seeded on collagen IV (400 μg/mL) and fibronectin (100 μg/mL) coated plates and cultured in EC media supplemented with 1X B27 and 10µM ROCK Inhibitor.

#### Immunofluorescence Staining

hPSC-BMELCs were seeded onto each slide of the Thermo Scientific™ Nunc™ Lab-Tek™ II Chamber Slide™ System (Thermofisher, 12-565-7). The cells were allowed to grow for 24 hours and were subsequently fixed with ice cold methanol for 15 minutes. The cells were then incubated with blocking buffer (10% Fetal Bovine Serum and 0.3% Triton-X 100 in PBS) for 30 minutes at room temperature. Primary antibodies were diluted in blocking buffer and incubated at 4°C overnight. After two washes with PBS, the cells were incubated with secondary antibody diluted in blocking buffer for 1 hour at room temperature. The chambers were washed twice with PBS and were stained with 1X DAPI for 3 minutes and mounted with coverslips using Fluoromount-G^TM^ Mounting Medium (Thermofisher, 00-4958-02). All slides were imaged using the Zeiss Axio-Observer 1 microscope and processed using ImageJ software.

#### Flow Cytometry

Cell media was aspirated, and cells were immediately lifted with either 200µL of StemPro^TM^ Accutase (for hPSC-BMELCs) or 200µL of Trypsin (all other cells). After 5-10 minutes of incubation at 37°C, the lifted cells were moved into a 96 well U-bottom plate and centrifuged at 1500rpm to pellet the cells. The cells were then resuspended in 200µL fixation buffer (2% Paraformaldehyde and 1% FBS in PBS) and fixed at 4°C for a minimum of 30 minutes. Samples were measured using the Miltenyi Biotec MACSQuant Analyzer 10 Flow Cytometer.

#### Plaque Forming Assay

All virus titers quantified as plaque forming units (PFU) in this study were done using BHK-21 cells cultured in MEM with 7.5% FBS. BHK-21 cells were seeded at high confluency (3.5×10^5^ cells/mL) in 6 well plates and were infected with 200µL virus inoculum at 24 hours post seeding. Virus inoculum was made by performing 1:10 serial dilutions of virus-containing sample in virus binding buffer (1%FBS in PBS). After virus addition, cells were gently agitated every 15 minutes for 1 hour for adsorption. The cells were then overlayed with 2.25% Avicel, 10% FBS, 1X Penicillin/Streptomycin, and 1X MEM NEAA in water and incubated at 37°C for 24 hours. The plates were then fixed with 4% paraformaldehyde for a minimum of 30 minutes at room temperature and then stained with crystal violet for a minimum of 30 minutes at room temperature. Finally, the plates were washed with water and left to dry overnight before plaques were counted.

#### Measurement of transendothelial electrical resistance (TEER)

The EVOM3 (World Precision Instruments) was used to measure TEER of hPSC-BMELCs in a transwell system. Specifically, the STX2 Plus electrodes were placed directly on top of the well such that each prong was in either the apical or basolateral chamber. The electrodes were placed at a depth that allows coverage of each electrode with media (approximately 0.5cm). The resistance values were typically measured beginning 1 day after seeding cells, along with a cell-free chamber control with plain media that was subtracted from the experimental wells. The resistance values (Ω) were multiplied by the surface growth area (cm^2^) to generate TEER (Ωcm^2^) values of the barrier formed by hPSC-BMELCs.

#### SVN and R47 poly-glycoprotein myc-E2 construct generation for VSV production

Generation of myc-tagged E2 using Hi-Fi Assembly have been previously described^69^. New England Biolabs (NEB) Hi-Fi Assembly primers were generated for SVN and R47 using the NEBuilder Assembly Tool software provided by NEB. Briefly, one set of primers was generated to amplify the SINV E3 gene while having overlapping sequences with the p278 pcDNA vector at the 5’ end and generating a myc tag at the 3’ end. An additional serine was added at the start of the myc sequence to allow for proper polyprotein cleavage. Another set of primers was generated to amplify SINV structural genes from E2 to E1 while adding a myc tag sequence at the 5’ end of the E2 sequence to overlap with the E3 fragment myc sequence and adding an overlapping sequence at the 3’ end complementary to the p278 pcDNA vector. These two fragments were generated using NEB’s Q5 Hi-Fidelity PCR protocol, and the vector was digested using NheI and ApaI. The three fragments were ligated together following NEB’s Hi-Fi Assembly kit, followed by bacterial transformation and construct amplification. The construct was verified using full plasmid sequencing services from Plasmidsaurus.

#### VSV Particle Production and infection

VSV particles pseudotyped with SVN or R47 glycoproteins were generated using previously published methods^43^. Briefly, HEK293T cells were seeded at a density of 2×10^5^ cells/mL in a 10cm dish with DMEM supplemented with 10% FBS. Cells were transfected with either SVN or R47 glycoprotein constructs the following day. One day after transfection, the cells were infected using VSV-ΔG seed particles and washed several times with PBS before cells were switched to DMEM supplemented with 2% FBS. The cells were infected for 24 hours before the supernatant was harvested. The infectious particle titer was calculated by performing a focus forming assay in Vero cells. Cells were infected by aspirating the medium, adding a low volume of virus inoculum, and incubated at 37°C with gentle agitation every 15 minutes for one hour. The inoculum was then aspirated, and cells were supplemented with the appropriate medium. Virus infection was determined via luciferase activity following the manufacturer’s instructions of the Promega Luciferase Assay System. Cell lysates were supplemented with luciferase substrate and read on a BioTek plate reader.

#### Fusion Assay

This assay was adapted from previous publications^44,45^. In short, hPSC-BMELCs were treated with bafilomycin A_1_ 1 hour prior to, 30 minutes prior to, at the same time as, 30 minutes after, or 1 hour after virus addition. For the time points prior to virus addition, hPSC-BMELC media was changed to media containing bafilomycin A_1_ at a concentration of 25nM. For all time points afterwards, bafilomycin A_1_ was added to the inoculum, such that the final concentration was 25nM. Five minutes prior to virus addition, cells were chilled on ice. The cells were kept at 4°C for one hour after virus addition. After the final treatment with bafilomycin A_1_, the cells were incubated at 4°C for an additional hour. The cells were washed twice with ice cold PBS, and media was replenished without bafilomycin A_1_, and moved to 37°C. Cells were infected at an MOI of 1 and incubated at 37°C for 12 hours when measuring viral RNA via RT-qPCR. The cells were infected at an MOI of 0.1 and incubated at 37°C for 18 hours before viral replication was measured via flow cytometry.

#### Virus Attachment and Internalization Assay

Alphavirus attachment and internalization were assessed according to previous publications^34^. Briefly, alphaviruses were diluted to varying MOIs in ice-cold 1%FBS in PBS. Five minutes prior to virus addition, cells were placed on ice. The cells were then inoculated with virus and were incubated for 90 minutes on ice and at 4°C. Upon completion of the incubation, the cells were washed 6 times with ice-cold PBS. For attachment studies, the cells were immediately lysed with TRIzol following the sixth wash. For internalization, the cells were replenished with media containing 25nM bafilomycin A_1_ and were allowed to incubate at 37°C for 1 hour. Cells were then chilled on ice and bound, uninternalized virions were digested with 0.5 mg/mL of proteinase K (Sigma, P2308) in PBS on ice for 2 hours. The cells were then washed an additional 6 times with ice-cold PBS, and were lysed with TRIzol for subsequent RNA extraction.

#### Quantitative Reverse Transcription Polymerase Chain Reaction (RT-qPCR)

Samples were washed once with PBS prior to lysis in TRIzol. Total sample RNA was isolated using the Direct-zol RNA miniprep kit (Zymo, R2050), and 600ng of total RNA was used to synthesize cDNA using Superscript III First-Strand Synthesis System (New England Biolabs) with random hexamer primers. The RT-PCR was set up by mixing diluted cDNA (1:1 0 in water) with Luna® Universal qPCR Master Mix (NEB, M3003X) and virus- or gene-specific primers and measured on a Bio-Rad CFX Opus96 RT-PCR System. For analysis, the housekeeping gene RPS11 cycle threashold values (Cq) values were first subtracted from the target gene Cq (ΔCq). The average ΔCq value of the control sample was then subtracted by the ΔCq values of the experimental samples resulting in the ΔΔCq value.

#### RNA isolation for RNA-Seq Analysis

Independent cultures of hPSC-BMELCs and HUVECs were established in 24-well plates for RNA extraction and submission to Novogene Corporation Inc. (California, USA) for whole RNA sequencing. Three wells were collected into one sample and triplicates of samples were submitted for RNA sequencing (n=3 per condition). For hPSC-BMELCs, each well contained 8e5 cells, for HUVECs, each well contained 5e5 cells. 24 hours after culture, cell pellets were collected and frozen at -80C for subsequent RNA extraction. RNA was extracted and DNAse-treated using the Qiagen mini RNA preparation kit (Maryland, USA) according to the manufacturer’s instructions. RNA concentration and quality was determined to match the guidelines of Novogene Corporation Inc. for RNA sequencing, concentration ≥20 ng/μL and RIN ≥7. Final libraries were generated from total RNA using KAPA mRNA Hyper kit, quantified, PolyA selected and sequenced non-strand specific using paired-end 150 bp (PE150) through Illumina PE150 technology. The data output was 20M read pairs (40M raw reads).

#### RNA-Seq Bioinformatic Analysis

The RNA-sequencing data were first processed to remove adapter sequences and low-quality reads followed by the alignment to the human genome GCHr38 using STAR (v2.6+)^70^. Count data from HTSeq were calculated^71^ and normalized using DESeq2’s median of ratios method (Ref. 3 - below). Afterwards, we used the normalized counts to run a differential expression analysis between the groups of interest using the DESeq2 R package^72^. In addition, R packages were used for various graphical visualizations, including ggplot2^73^ for PCA plot and pheatmap (R package version 1.0.12) for heatmap^74^. Finally, the Database for Annotation, Visualization and Integrated Discovery (DAVID; version 6.8) was used to perform gene ontology (GO), functional annotation and functional annotation clustering of differentially expressed genes^75^. A false discovery rate (FDR) of 0.05 was selected as the cut-off criterion for functional annotation. The classification stringency setting used for functional annotation clustering was medium with default setting for function grouping, except for EASE which was lowered to 0.00003 to reduce inclusion of non-significant terms into the clusters. Annotation Clusters of significantly overrepresented groups with terms having a FDR < 5% were accepted for further consideration.

#### Antibody Blockade

Alphavirus receptor blockade was adapted from previous work^32^. In short, HEK293T cells were seeded at a density of 4×10^5^ cells/mL in a 24 well plate. The following day, cells were incubated with 100µL of 100µg/mL antibody for 1 hour at 37°C prior to infection. Following 1-hour incubation, 50µL of virus was added to the cells such that each well was infected at an MOI of 1. The virus was allowed to adsorb for an additional hour at 37°C. The inoculum was then fully aspirated, and the cells were supplemented with DMEM with 10% FBS. The cells were infected for 18 hours and then processed for flow cytometry analysis.

#### Proteomics sample processing

The Pierce™ Cell Surface Biotinylation and Isolation Kit (Thermofisher, A44390) was appended with the EasyPep™ MS Sample Prep Kit (Thermofisher, A40006) to generate the cell surface proteomics dataset according to manufacturer’s protocols. Briefly, cells were detached from the culture vessel and washed 5 times with DPBS, followed by biotinylation of outer proteins using Sulfo-NHS-SS-Biotin (Thermofisher). Cells were then lysed using the provided lysis buffer, and biotinylated proteins were enriched using neutravidin beads. The isolated biotinylated proteins were then reduced, alkylated, and trypsinized using Trypsin/Lys-C, followed by peptide clean-up using columns provided by the EasyPep™ MS Sample Prep Kit.

#### Mass spectrometry proteomics acquisition

Dried peptides were resuspended in 0.1% (v/v) Formic acid (FA, LC/MS grade, Sigma Aldrich) in water (LC/MS grade, Fisher Scientific) and analyzed on a timsTOF HT mass spectrometer (Bruker Daltonics), paired with a Vanquish Neo ultra-high pressure liquid chromatography system (Thermo Fisher Scientific). Mobile phase A consisted of 0.1% (v/v) FA in water (LC/MS grade, Fisher Scientific), and mobile phase B consisted of 0.1% (v/v) FA in 100% Acetonitrile (LC/MS grade, Fisher Scientific). The LC was operated in trap-and-elute mode, where the peptides were first trapped onto a PepMap Neo Trap column (5 mm, 100 Å pore size, 5 µm particle size) and then reversed-phase separated using gradient mentioned below on a Bruker PepSep C18 reverse phase column (15 cm, 150 µm I.D., 100 Å pore size, 1.5 µm particle size), kept at 50°C using a column oven for Bruker CaptiveSpray source (Sonation Lab Solutions). The capillary voltage was set to 1700 V. The peptides were separated at 500 nL/min flow rate, on a gradient of %B as follows: 5% to 50% B over 40.5 min, then to 60% in the next 1 min, followed by a column wash step where %B was increased to 95% B in 0.5 min, and stayed same for next 3 min at 1.5 µL/min flow rate. The raw data was acquired in data-independent acquisition coupled with parallel accumulation–serial fragmentation (dia-PASEF) mode. Using equal-size windows of 25 Da with an overlap of 1 Da in the m/z *vs* ion mobility plane, the precursor ion coverage was maximized for further MS/MS. For dia-PASEF, in the ion mobility (1/K0) range 0.64 to 1.40 Vs cm-2, the collision energy was linearly decreased from 59 eV at 1/K0 = 1.40 Vs cm-2 to 20 eV at 1/K0 = 0.64 Vs cm-2 to collect the MS/MS spectra in the mass range 254.2 to 1287.2 Da. The estimated mean cycle time was 1.59s. The ion accumulation time and ramp times in the dual TIMS analyzer were set to 100 ms each.

#### Mass spectrometry proteomics data analysis

The raw files were processed with DIA-NN ver. 1.9.2 in two steps. The first step generated an *in-silico* spectral library using reviewed protein entries from the UniProt Human database (downloaded Nov. 11, 2024) and the DIA-NN default common contaminants list. For precursor ion generation, ‘FASTA digest for library free search/library generation’ and ‘Deep learning-based spectra, RTs, and IMs prediction’ options were selected. Trypsin/P was selected as protease, with a maximum of two missed cleavage and three variable modifications allowed. Carbamidomethylation at Cysteine was kept as a fixed modification. The variable modifications used were: N-term M excision, Oxidation (M), Acetylation (N-term), and CAMthiopropanoyl (K, N-term, Unimod Accession # 293). Other parameters used for the library generation are- Peptide length range: 7 to 30, Precursor charge range: 1-4, Precursor m/z range: 250-1800, and Fragment ion m/z range: 200-1800. The generated spectral library was used to search the raw files. For the DIA-NN search, both MS1 and MS2 mass accuracy was set to automatic inference, with the match-between-run option selected. Other search parameters were set to default settings.

Identified peptides were next evaluated for quality and replicate consistency. First, an abundance correlation matrix of peptide intensities and identities was generated to evaluate the correlation between replicates. Samples C+.2 and iB+.2 were excluded due to poor correlation between replicates. Next, based on the number of unique peptides and proteins detected, sample C-.1 was excluded because it contained a substantially higher number of peptides and proteins compared to its replicates, and sample U-.2 was excluded due to an insufficient number of peptide and protein identifications. After this quality control step, the DIA-NN report.tsv output file was imported into MSstats for cross-run normalization, protein summarization, and statistical comparative analysis between conditions. MSstats performs normalization by median equalization, imputation of missing values and median smoothing to combine intensities for multiple peptide ions or fragments into a single intensity for their protein group, and statistical tests of differences in intensity between infected and control time points. Default settings for MSstats were used for adjusted P values. By default, MSstats uses the Student’s t-test for P value calculation and the Benjamini–Hochberg method of FDR estimation to adjust P values. To identify the background proteins (nonspecific binders), we compared proteins detected in the + biotin and - biotin samples for each cell line. Proteins with an adjusted p-value < 0.05 and log2 fold change > 1 in the + biotin vs. – biotin comparison were identified as foreground proteins, specific to the biotin labeling, for each cell line. Using the list of foreground proteins only, we next compared the + biotin samples across the three cell lines in the + biotin conditions to identify differences in their cell surface proteomes. Proteins not included in the foreground list were excluded to ensure specificity to the enriched proteome.

#### Western Blotting

Samples were lysed in homemade RIPA Buffer (150mM NaCl, 1% Nonidet P-40 Subtitute, 0.5% Sodium Deoxycholate, 0.1% SDS, 50mM Tris-HCl, pH 7.4) with Roche cOmplete Mini, EDTA-free Protease Inhibitor Cocktail (Millipore Sigma, 11836170001). Lysates were clarified at >16,000rpm for 5 minutes, and protein concentration was measured using Pierce^TM^ BCA Protein Assay Kit (ThermoFisher, 23225) according to manufacturer’s protocols. Samples were denatured by mixing 20µg of protein with 4x Laemmli buffer and boiled at 100°C for 10 minutes. The samples were separated by 12%–15% sodium dodecyl sulphate–polyacrylamide gel electrophoresis (SDS-PAGE) and transferred onto methanol-activated 0.2 μm PVDF membranes. The membranes were blocked with 5% milk in PBS-T (0.1% Tween-20 in PBS) for 30 minutes at room temperature. Primary antibodies were diluted in 5% milk in PBS-T and incubated with the membranes overnight at 4°C. After three washes with PBS-T, the membranes were incubated with secondary antibodies for at least 1 hour at room temperature. Finally, a chemiluminescent HRP substrate was applied to the membrane for detection of proteins. Protein detection was done using a ChemiDoc Imager (Bio-Rad), and images were processed using ImageLab (Bio-Rad).

#### Quantification and statistical analysis

Image analyses and quantification were performed using ImageJ or ImageLab. Graphpad Prism 10 was used for all statistical analysis. Results are shown as means ± standard deviation (SD) unless stated otherwise. All data shown are representative of at least 2 independent experiments. Each independent experiment has a minimum of 3 biological replicates per condition. Statistical significance was indicated as: all ns (no significance) are not shown, ^∗^ (p < 0.05), ^∗∗^ (p < 0.01), ^∗∗∗^ (p < 0.001) and ^∗∗∗∗^p ≤ 0.0001 in the figures and figure legends unless stated otherwise.

## FIGURE LEGENDS

**Supplemental Figure 1.**
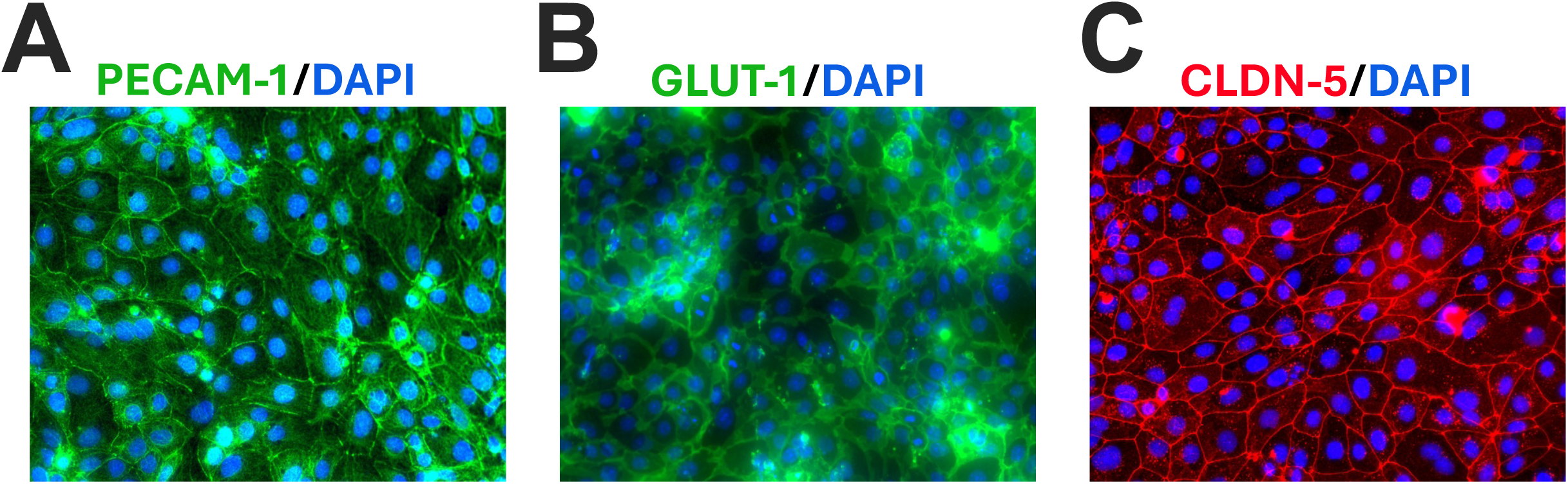
hPSC-BMELCs express appropriate BMEC-specific markers. hPSC-BMELCs were fixed with ice cold methanol 24 hours after seeding and stained for platelet endothelial cell adhesion molecule 1 (PECAM-1, also named CD31) (A), glucose transporter 1 (GLUT1) (B), or claudin 5 (CLDN-5) (C). One representative image is shown from 3 independent experiments.

**Suppemental Figure 2.**
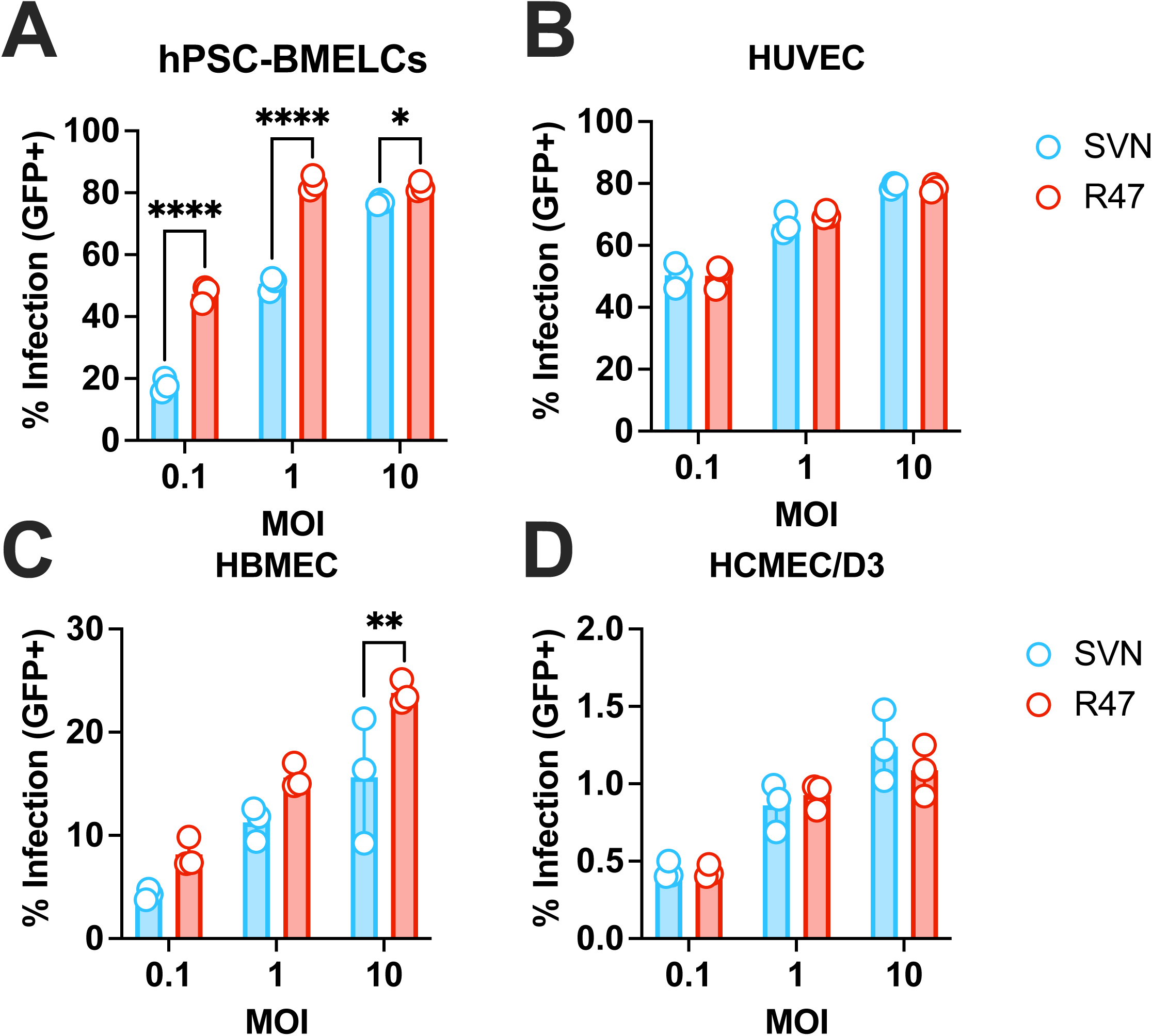
SINV infects endothelial cell models to varying degrees. GFP-expressing SVN and R47 were used to infect hPSC-BMELCs (A), HUVECs (B), primary BMECs (C), or immortalized BMECs (D) at MOIs of 0.1, 1, or 10. Cells were harvested 24 hours post infection for flow cytometry analysis. Asterisks indicate statistically significant differences in infection between SVN and R47 at the indicated MOI (two-way ANOVA and Šidák’s multiple comparisons): *, *p* < 0.05; **, *p* < 0.01; ****, *p* < 0.0001. Data are representative of 2 independent experiments. Data is shown as mean ±SD.

**Supplemental Figure 3.**
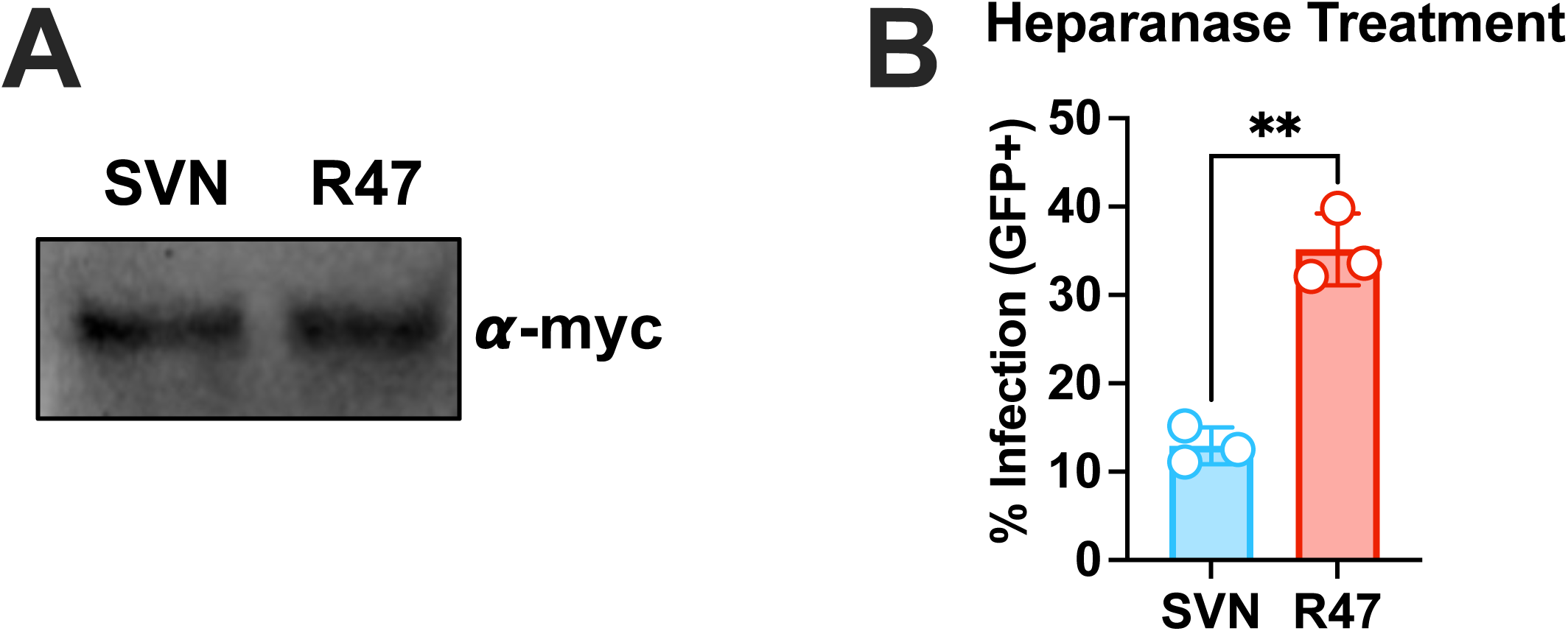
Differences in infection with viruses carrying SVN vs R47 glycoproteins are independent of unequal E2 distribution on VSV particles and independent of heparan sulfate binding. (A) Pseudotyped VSV particles were lysed in SDS and normalized by protein concentration. Samples were separated by PAGE and N-terminally myc tagged E2 was probed using anti-myc. One representative blot is shown from two independent VSV preparations. (B) hPSC-BMELCs were pretreated with heparanases I and III for 1 hour at 37°C prior to infection with GFP-expressing SVN or R47. Cells were infected at an MOI of 0.1 and harvested 24 hours post infection. Cells were analyzed via flow cytometry. Asterisks indicate statistically significant differences between SVN and R47 infection (unpaired t-test): **, *p* < 0.01. Data are representative of 2 independent experiments. Data is shown as mean ±SD.

**Supplemental Figure 4.**
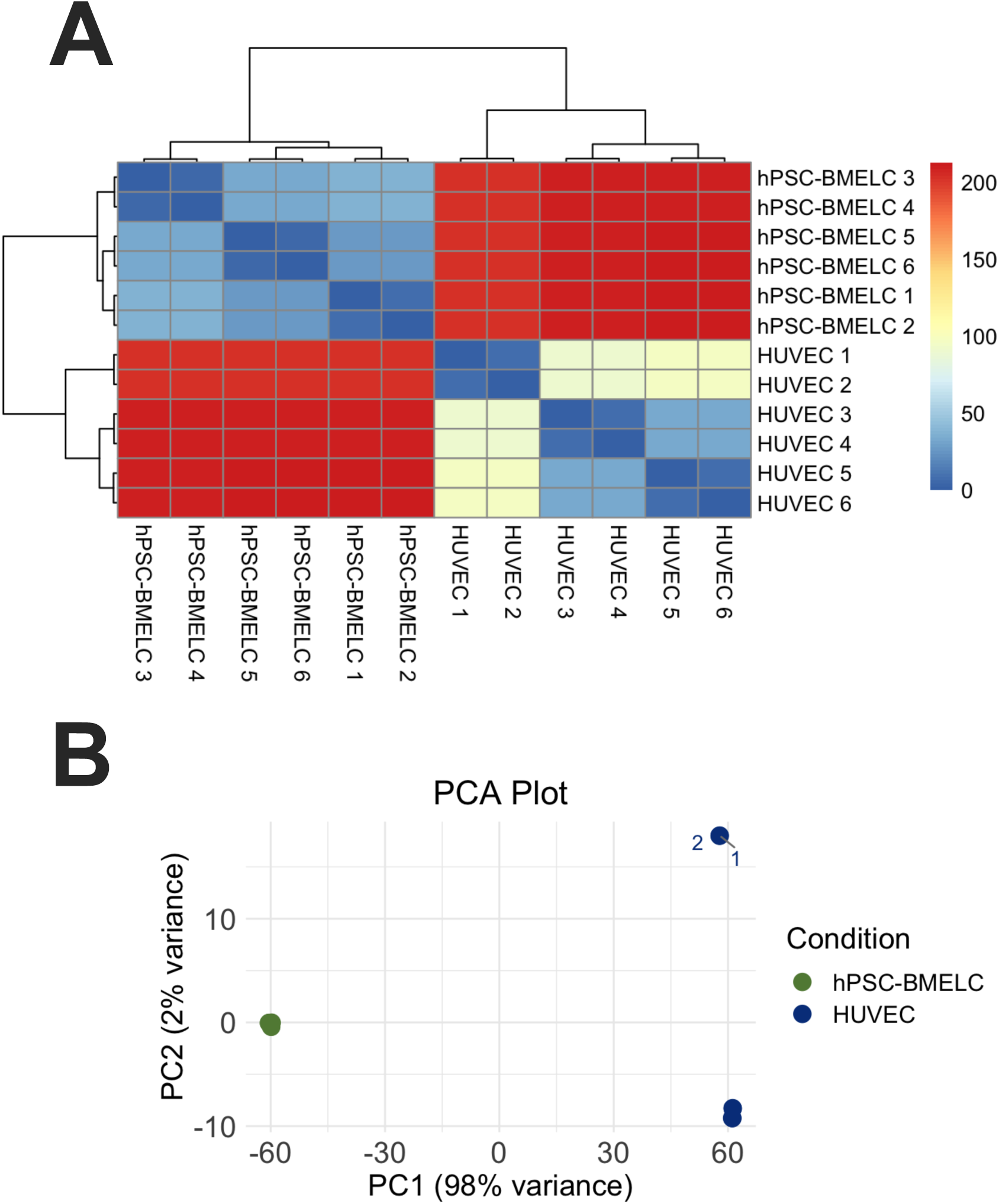
hPSC-BMELCs and HUVECs have distinct gene expression profiles. RNA was isolated from hPSC-BMELCs and HUVECs under basal conditions. (A) Sample to sample heatmaps were generated to compare variability across hPSC-BMELC and HUVEC samples. Heatmaps were generated in R using the pheatmap and ggplot packages. (B) Principal component analysis (PCA) showing the differences across all samples, with the difference between hPSC-BMELC and HUVEC samples explaining 98% of the total variance, while two HUVEC samples that cluster away from the rest of the HUVEC samples only explain 2% of the total variance. Transcriptomic analysis and read normalization were generated by DESeq2 analysis on R.

**Supplemental Figure 5.**
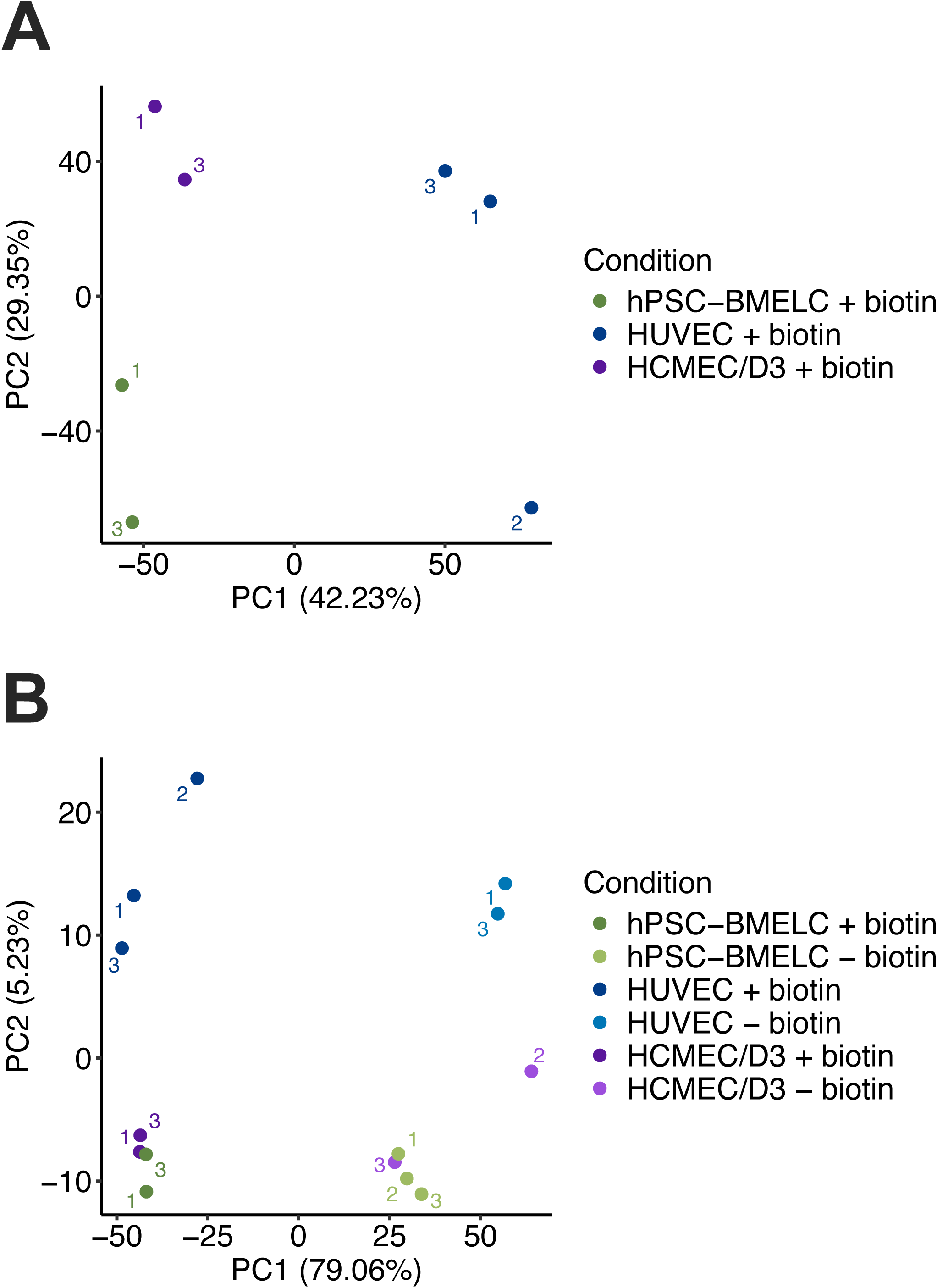
hPSC-BMELCs share characteristics with other endothelial cells and biotinylated sample profiles are distinct from non-biotinylated samples. (A) Principal component analysis of biotinylated samples from all three cell types: hPSC-BMELCs (green), HUVECs (blue), HCMEC/D3 (purple). (B) Principal component analysis of biotinylated (darker shade) and non-biotinylated (lighter shade) samples from all three cell types: hPSC-BMELCs (green), HUVECs (blue), HCMEC/D3 (purple).

## Notes

https://www.ncbi.nlm.nih.gov/geo/query/acc.cgi?acc=GSE287658

### Competing Interest Statement

The authors have declared no competing interest.

